# Cortical excitability controls the strength of mental imagery

**DOI:** 10.1101/093690

**Authors:** Rebecca Keogh, Johanna Bergmann, Joel Pearson

## Abstract

Mental imagery provides an essential simulation tool for remembering the past and planning the future, with its strength affecting both cognition and mental health. Research suggests that neural activity spanning prefrontal, parietal, temporal, and visual areas supports the generation of mental images. Exactly how this network controls the strength of visual imagery remains unknown. Here, brain imaging and transcranial magnetic phosphene data show that lower resting activity and excitability levels in early visual cortex (V1-V3) predict stronger sensory imagery. Electrically decreasing visual cortex excitability using tDCS increases imagery strength, demonstrating a causative role of visual cortex excitability in controlling visual imagery. These data suggest a neurophysiological mechanism of cortical excitability involved in controlling the strength of mental images.

## Introduction

Visual imagery - the ability to “see with the minds eye” - is ubiquitous in daily life for many people, however the strength and vividness with which people are able to imagine varies substantially from one individual to another. Due to its highly personal nature, the study of visual imagery has historically relied on self-report measures and had long been relegated to the shadows of scientific inquiry. However, with the advent of fMRI and new analysis techniques like decoding, as well as new advances in behavioral and psychophysical experiments, this is quickly changing (*1*). Despite these advances, very little research has investigated why such large individual differences in the ability to imagine exist. Much of the past research focused on the similarities between visual imagery and perception, and has shown that a large network of occipital, parietal, and frontal areas are involved when imagining (*2*). Here, we used a multi-method approach (fMRI, TMS and tDCS) to assess the potential contributions of resting levels of cortical excitability in the visual imagery network as a critical physiological precondition, which determines the strength of visual imagery.

To measure mental imagery strength, we utilized the binocular rivalry imagery paradigm, which has been shown to reliably measure the sensory strength of mental imagery through its impact on subsequent binocular rivalry perception(*1, 3*). Previous work has demonstrated that when individuals imagine a pattern or are shown a weak perceptual version of a pattern, they are more likely to see that pattern in a subsequent brief binocular rivalry display, see (*2*) for review of methods). Longer periods of imagery generation, or weak perceptual presentation, increase the probability of perceptual priming of subsequent rivalry. For this reason, the degree of imagery priming has been taken as a measure of the sensory strength of mental imagery. Importantly, this measure of imagery is directly sensory; while it is related to subjective reports of imagery vividness, it is not a proxy for metacognitive reports of imagery vividness, and findings regarding their relationship across individuals have been mixed (see supplementary figure S1A and (*4, 5*)). This measure of imagery strength has been shown to be both retinotopic location- and spatial orientation-specific (*5, 6*), is reliable when assessed over days or weeks (see supplementary figure S2A and (*5*)), is contingent on the imagery generation period (therefore not due to any rivalry control) and can be dissociated from visual attention (*6*). This measure of imagery is advantageous in that it allows us to avoid the prior limitations of subjective introspections and reports, which can often be unreliable and swayed by the context and an individual’s ability to introspect (recently referred to as metacognition). Additionally, metacognition appears to be dissociable from the imagery itself, with training-based improvements in imagery metacognition occurring without changes in imagery strength (*7*).

To measure cortical excitability and it’s role in visual imagery strength fMRI and TMS were used, and non-invasive brain stimulation (in the form on transcranial direct current stimulation (tDCS)) was employed to manipulate cortical excitability.

## Results

### Correlations between visual cortex excitability and visual imagery strength

For a first assessment of the relationship between cortex physiology and imagery strength, we looked at fMRI data we assessed a sample of 31 participants resting-state data (these participants have previously been reported upon in (*5, 8*) however these analyses were structural rather than functional). We related this data set to each individual’s imagery strength determined using the binocular rivalry method (% primed, see Fig.1A). Using a whole-brain surface-based group analysis (see Methods), we found that the normalized mean fMRI intensity of clusters in the visual cortex showed a negative relationship with imagery strength, while frontal cortex clusters showed positive relationships (multiple comparison-corrected; see Fig. S3 and Supplementary Table S1 and S2).

**Figure. 1.**
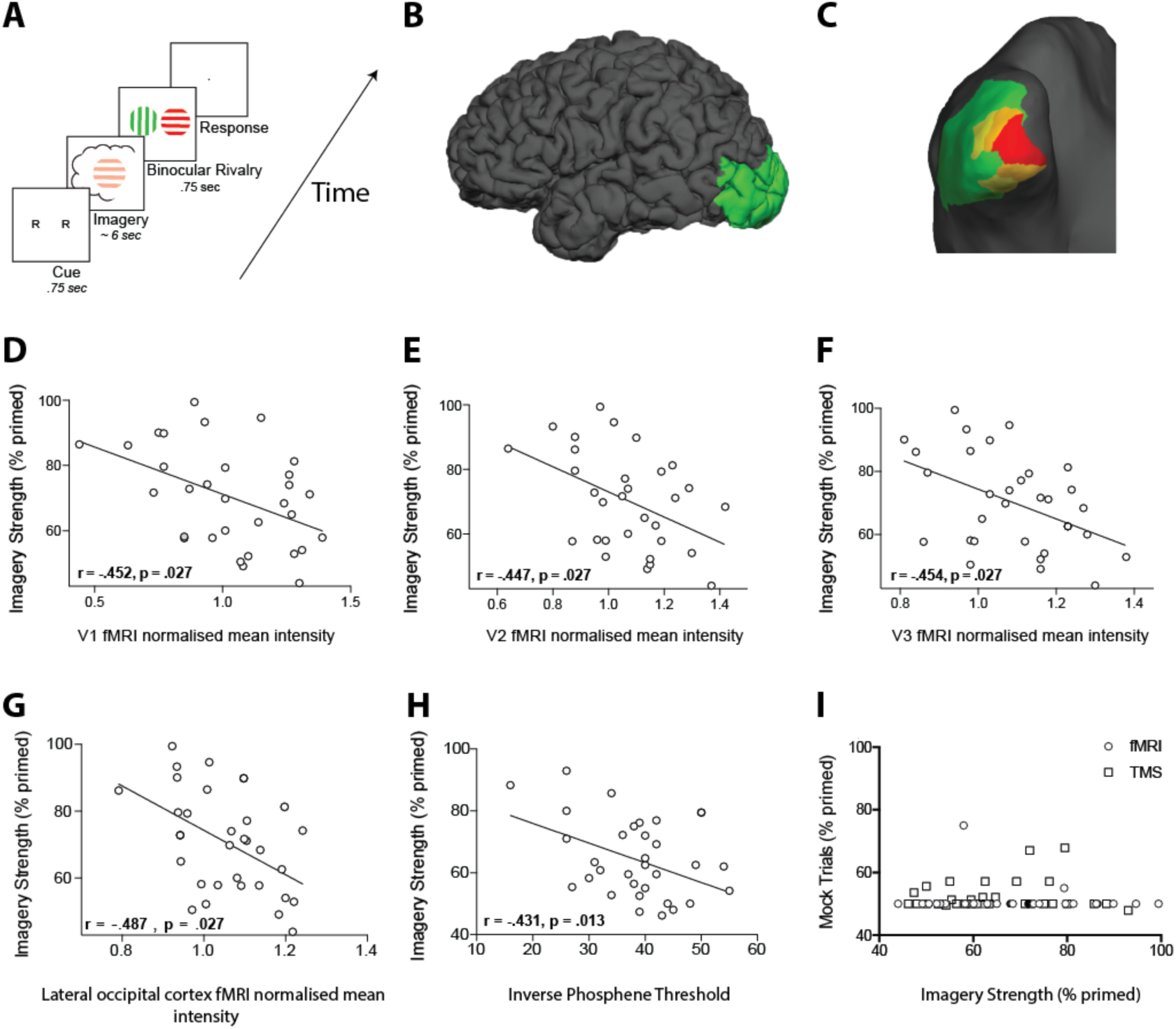
A. Timeline of the basic imagery experiment. Participants were cued to imagine a red-horizontal or a green-vertical Gabor patch for six to seven seconds by the letter R or G (respectively). Following this, they were presented with a brief binocular rivalry display (750ms) and asked to indicate which image was dominant. In the behavioral experiments with the brain-imaging sample and in three of the tDCS experiments, a rating of subjective vividness of the imagery also preceded the binocular rivalry display. **B and C.** Lateral view of the pial surface (B) and posterior view of the inflated surface (C) of the visual areas that showed a significant negative relationship with imagery. Red = V1, Orange = V2, Yellow = V3 and Green = lateral occipital area **D-G.** Correlation between normalized mean fMRI intensity levels in V1, V2, V3 and lateral occipital area and imagery strength. Individuals with lower mean fMRI intensity levels in early visual cortex showed stronger imagery. **H.** Correlation between the inverse phosphene threshold and imagery strength. Individuals with lower cortical excitability in visual cortex tended to have stronger imagery. **I.** Correlation between mock priming scores and real binocular rivalry priming for participants in the fMRI (circles) and TMS (squares) study. There was no significant association between perceptual priming in real and mock trials for the fMRI or TMS data. In the scatterplots (D-I), each data point indicates the value of one participant; the bivariate correlation coefficients are included with their respective significance levels.

To further investigate these relationships, we first focused on the visual cortex. We mapped early visual areas V1, V2 and V3 (estimated with standard fMRI retinotopic mapping; see Methods) and the adjacent occipito-parietal areas (defined by the Desikan–Killiany atlas). Following this, we related the normalized mean fMRI intensity scores of each area to each participant’s imagery strength; four ROIs showed a significant negative relationship with imagery strength (V1: *r*= -.45, *p* = .027; V2: *r*= -.45, *p* = .027; V3 *r*= -.45, *p* = .027; lateral occipital area, r= -.49, *p* = .027; FDR-adjusted p-values to correct for multiple comparisons: Fig.1D-G).

Despite prior work demonstrating that the binocular rivalry measure of imagery is specific to prior imagery generation and not attentional control of the subsequent rivalry presentation, we utilised the known perturbative effect of bright background luminance on imagery generation (*6, 9, 10*). Twenty-two of the same participants had completed the same imagery task again, however this time with uniform and passive background luminance during the seven second imagery generation. We hypothesized that if high background luminance perturbs imagery, then the relationship between imagery under high-luminance conditions and fMRI intensity should be reduced. As predicted, imagery strength measured in the presence of background luminance did not significantly correlate with the normalized mean fMRI intensity measure for any visual area (V1: r = -.18, *p* = .41, V2: r = -.32, *p* = .15, V3: r = -.40, *p* = .07, lateral occipital cortex: r = -.34, p = .12). For V1 and V2 these correlations were significantly smaller than their corresponding ‘no-luminance’ correlations (one-sided Steiger’s Z-test, V1: Z(19) = −1.83, *p* = .032, V2: Z(19) = −1.77, *p* = .035, multiple comparison-corrected), however the difference in the correlations for V3 and lateral occipital area were not (V3: Z(19) = −1.57, *p* = .065, lateral occipital area Z(19)=-1.00, *p* = .157, multiple comparison-corrected). As the luminance never co-occurred with the rivalry presentation, it should only interfere with the generation of the images and not the attentional or volitional control of rivalry, suggesting the physiology-behavior relationship cannot be explained by voluntary control of binocular rivalry.

Our data are compatible with the hypothesis that the resting levels of early visual cortex activity are negatively related to imagery strength. However, mean fMRI intensity levels are not a commonly used measure. Individual variation in this parameter is influenced by many factors that are challenging to control for, e.g. differences in proton density(*11*) . Additionally, fMRI activity at rest is influenced by a number of factors other than neural activity, such as non-neuronal physiological fluctuations or scanner noise (*12*). Nonetheless, previous research has shown that the fMRI signal during resting state is strongly reflective of underlying neural activity (*13, 14*). By normalizing the individual fMRI signal levels of our ROIs using the whole brain’s signal intensity, we aimed to control for some non-neuronal influences that may affect the individual brain in its entirety (e.g. scanner noise). In addition, further analyses of head motion indicate that it is unlikely that individual differences here had an influence on the pattern of significant relationships (see Methods and Supplementary Results).

We also excluded the possibility that differences in head size contributed to the relationship between the ROIs’ mean fMRI signal levels and individual behavior. Using cortical surface area and volume, respectively, as proxies for head size there was no significant relationships with either the behavioral or fMRI data (all *p*>.16). Accordingly, partialling out these factors did not change the pattern of significant results (all *p* < .02). Furthermore, a surface-based whole-brain group analysis of another fMRI resting-state measurement, which included almost all of the previous participants, indicates that the relationships are reliable (see Supplementary Figure S3).

To further substantiate our observations and circumvent other potential confounds that might influence the fMRI data, we next utilized a different methodology that measures cortical excitability: transcranial magnetically induced phosphenes. A new sample of 32 participants performed an automated phosphene threshold (PT) procedure using transcranial magnetic stimulation (TMS) over early visual cortex (see methods). Visual phosphenes are weak hallucinations caused by TMS applied to visual cortex. The magnetic strength needed to induce a phosphene is a reliable and non-invasive method to measure cortical excitability. In line with the normalized mean fMRI intensity data, we found a significant negative correlation between imagery strength and visual cortex excitability (data shows inverse phosphene threshold (100-PT) for easy of visualizing data as PT’s are negatively correlated with cortical excitability: *r*= -.44, *p* = .0127; Fig.1H). In other words, individuals with lower visual cortex excitability exhibited stronger imagery. Importantly, we also tested the phosphene threshold retest reliability for our paradigm over two days and found it was a very reliable measure (*r_s_* = .75, *p* < .001; see Supplementary Figure S2B) and the imagery strength re-test was also reliable (tDCS experiments: *r* _s_= .51, *p* < .0001; see Supplementary Figure S2A).

To assess possible effects of a decisional bias, mock rivalry trials were included in all tests of imagery strength (*6, 9, 10, 15*)(see Methods). We found no correlation between real binocular rivalry and ‘mock priming’ (fMRI (circles r_s_ = -.03, *p* = .89 & TMS (squares) *r_s_* = -.01, *p* = .97, see Fig.1I). These data, in conjunction with the effects of background luminance for the MRI data, make it unlikely that the relationship between imagery strength and physiology is due to demand characteristics, decisional bias or voluntary rivalry control.

### Manipulating visual cortex excitability

The data suggest that the excitability of the visual cortex plays a role in governing the strength of visual imagery, as participants with lower visual cortex activity tended to have stronger visual imagery and vice versa. However, these data do not speak to the causal role of early visual cortex in creating strong mental images. If the association between imagery strength and visual cortex activity is causal, manipulating visual cortex excitability should likewise modulate imagery strength.

To assess this hypothesis, we utilized non-invasive transcranial direct current stimulation (tDCS), which can increase or decrease cortical excitability depending on electrode polarity and position (see (*16*) for review). Broadly speaking when the cathode is placed over the cortex, the underlying excitability is decreased, whereas the anode increases excitability. Sixteen new participants underwent both anodal and cathodal stimulation of visual cortex (see Fig.2B for electrode montage) on two separate days (separated by at least twenty-four hours). The reference electrode was placed on the supraorbital cortex. On each day, participants completed six blocks of the imagery task, two before tDCS, two during tDCS and two post tDCS (see Fig. S4A for experimental timeline). To assess the effect of tDCS on imagery strength, we calculated the percent change in priming for each participant from baseline (one each day) by subtracting the mean priming rate preceding tDCS from those during and after tDCS, such that positive numbers indicate increases from baseline and negative ones indicate decreases. This was then divided by their baseline imagery strength, to control for an individual’s imagery strength, and multiplied by 100 ((block (n) - (average of two tDCS blocks before stimulation)/ (average of two tDCS blocks before stimulation))*100). These percent change scores were used as we were interested in how much each individual’s imagery strength increased or decreased relative to their baseline imagery scores for the different tDCS polarities. Analyzing data in this way also normalized the data for individual differences in visual imagery strength. Positive values indicate an increase in imagery strength, whereas negative values a decrease.

**Figure. 2.**
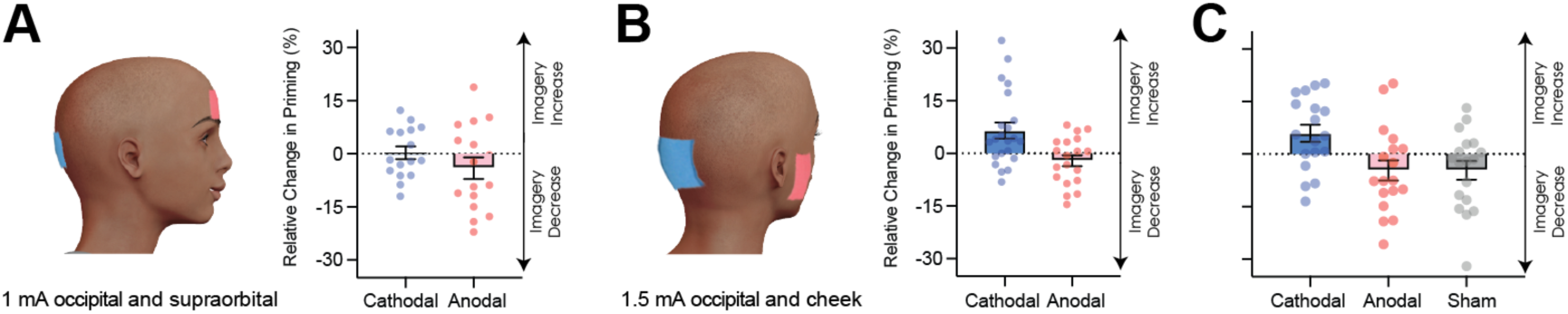
Visual cortex stimulation data. **A.** Effect of visual cortex stimulation on imagery strength at 1mA. The left image shows the tDCS montage, with the active electrode over Oz and the reference electrode on the supraorbital area. The right image shows the effect of cathodal (decreases excitability, blue dots represent each participant’s data) and anodal (increases excitability, red dots represent each individual participant’s data) stimulation averaged across all tDCS stimulation blocks (D1, D2, P1 and P2). **B.** Effect of visual cortex stimulation on imagery strength at 1.5mA. To the left: the tDCS montage with the active electrode over Oz and the reference electrode on the right cheek. To the right: the effect of cathodal (blue dots, decrease excitability) and anodal (red dots, increase excitability) stimulation averaged across all blocks during and after tDCS stimulation (D1, D2, P1 and P2). Each data point represents a single participant. Imagery strength increases in the cathodal stimulation condition (blue), when neural excitability is reduced. **C.** Effect of visual cortex stimulation on imagery strength at 1.5mA (same montage as in 2B). The left bar shows the relative change in imagery strength for cathodal stimulation (blue bar, blue dots represent individual participants data), the middle bar shows the relative change in imagery strength for anodal stimulation(red bar, red dots represent individual participants data), while the right bar shows the change in imagery strength for sham stimulation (grey bar, grey dots represent individual participants data). All error bars show ±SEMs.

Figure 2A shows relative imagery priming percent change scores averaged across all stimulation blocks with 1mA of tDCS stimulation (data per block can be seen in S4C) for anodal (red bars and data points) and cathodal (blue bars and data points) stimulation. Due to some missing data in the following tDCS experiments, and to control for the potential effect of stimulation order on our results, a linear mixed effects analysis was computed for all following experiments. This analysis was run with a 2 (tDCS polarity: cathodal and anodal), x 4 (block:D1, D2, P1, P2 – see S4A for timeline and S4C for data for each block) x 2 (order of stimulation: cathodal on first or second day) design. When fitting a linear mixed model the effect of tDCS polarity was not significant (χ^2^(1) = 2.99, *p* = .084).

The inconclusive results from the first tDCS experiment may be due to the stimulation intensity of 1 mA being too low to produce any effect (many tDCS studies use an intensity ranging from 1.5-2mA, for example see (*17*)). To investigate whether the lack of a significant result with 1mA was due to the intensity of the stimulation being too low, we ran a second tDCS study with a higher intensity of 1.5mA (see methods) and both cathodal (blue bars and dots) and anodal (red bars and dots) stimulation conditions. Additionally, to ensure we were not also stimulating the pre-frontal cortex, the supraorbital placement of the reference electrode was moved to the cheek (Fig.2C). A linear mixed effects analysis was run with a 2 (tDCS polarity:cathodal and anodal), x 4 (block:D1, D2, P1, P2 – see S4A for timeline and S4E for data for each block) x 2 (order of stimulation: cathodal on first or second day) design. The effect of tDCS polarity was significant χ^2^(1) = 15.85, *p* = 6.86e^-05^). The changes were in line with the correlational data for resting levels of visual cortex excitability and activity (see figure 1), such that participants imagery strength increased when visual cortex excitability was decreased (cathodal stimulation), while the opposite was true of increasing visual cortex excitability (anodal stimulation).

It is likely that the change from 1mA to 1.5mA allowed us to observe the modulatory effects of tDCS; however it also might be that the change in montage had an influence (i.e. location of reference electrode). It may also be the case that there are either fatigue or practice effects on this visual imagery task, i.e. perhaps participants just get better/worse on this task due to doing multiple sessions of the task. For this reason, a third experiment was run to assess the effects of fatigue/practice and the change of reference location. This study was identical to the above study with the inclusion of a sham condition where the tDCS machine shut off after thirty seconds of stimulation. A linear mixed effects analysis was run with a 3 (tDCS polarity: cathodal, anodal and sham), x 4 (block:D1, D2, P1, P2 – see S5A for timeline and S5C for data for each block) x 3 (order of stimulation: cathodal on first, second or third day) design. The effect of tDCS polarity was again significant (χ^2^(2) = 21.66, *p* = 1.98e^-05^). These data indicate that cathodal stimulation results in increased imagery strength (see figure 2C), and this is unlikely to be a practice effect, as sham stimulation results in decreases in imagery strength. Additionally previous work using the same binocular rivalry paradigm has demonstrated no increases in visual imagery strength from multiple days of training(*4*). Taken together these data suggest that cathodal stimulation leads to increases in imagery strength due to decreased visual cortex excitability, and these changes cannot be explained as a learning effect due to performing multiple sessions of the imagery task.

Although other studies have provided evidence that tDCS does change the excitability of the visual cortex (see (*18*) for example), we wanted to ensure that our specific stimulation paradigm was indeed modulating visual cortex excitability. We ran a separate control study comparing TMS-phosphene thresholds before and after the same tDCS paradigm (1.5mA, active electrode on Oz and reference on cheek, see Figure 2D, all subjects received both anodal and cathodal stimulation across separate days; see methods for further details). If our cathodal stimulation is decreasing visual cortex excitability, greater TMS power output would be required to elicit phosphenes post cathodal stimulation, whereas post anodal stimulation we would predict the opposite effect. A linear mixed effects analysis was run with a 2 (tDCS polarity: cathodal and anodal), x 2 (block: Pre tDCS and Post tDCS) x 2 (order of stimulation: cathodal on first or second day) design. We found that phosphene thresholds measured immediately after anodal stimulation decreased, whereas after cathodal stimulation phosphene thresholds increased (significant effect of tDCS polarity (χ^2^(1) = 4.3245, *p* = .0376, see figure S6)). These findings show that our stimulation paradigm changes cortical excitability in the expected direction, i.e. cathodal stimulation decreases cortical excitability, whereas anodal stimulation increases activity.

In summary we found that in two separate experiments resting levels of early visual cortex excitability/activity negatively predicted visual imagery strength (fMRI and TMS, figure 1D-H). We were also able to causally alter visual imagery strength in two separate tDCS experiments. Specifically decreasing visual cortex excitability (using cathodal stimulation 1.5mA) increased imagery strength (see figure 2B & C).

### Correlations between frontal cortex excitability and imagery strength

Our data suggest that visual cortex excitability plays a causal role in modulating imagery strength, but how exactly might excitability influence imagery strength? One hypothesis is that hyperexcitability might act as a source of noise in visual cortex that limits the availability or sensitivity of neuronal response to top-down imagery signals, resulting in weaker image-simulations. This hypothesis is supported by behavioral work showing that both imagery and visual working memory can be disrupted by the passive presence of uniform bottom-up afferent visual stimulation (*9, 10*), known to increase neural depolarization in primary visual cortex (*19*). However, the strength of the top-down imagery-signals arriving at visual cortex should also play a role in governing imagery strength as activity in a brain network including prefrontal areas supports mental image generation(*2*).

As mentioned previously, the multiple comparison-corrected whole-brain surface-based analysis of the mean fMRI intensity levels at rest revealed relationships with clusters in both visual cortex and frontal cortex (see Supplementary Fig. S3 and Table S2). Most of the significantly positive frontal clusters were located in superior frontal cortex. Additionally, using a ROI-based approach, normalized mean fMRI intensity levels in two frontal areas also showed positive relationships with imagery strength: superior frontal cortex (Fig.3A-B dark blue: *r* = .41, *p* = .022) and area parsopercularis (*r* = .38, *p* = .033; ROIs defined by the Desikan–Killiany atlas). However, these relationships did not survive multiple comparison correction (both *p* >.05).

**Figure. 3.**
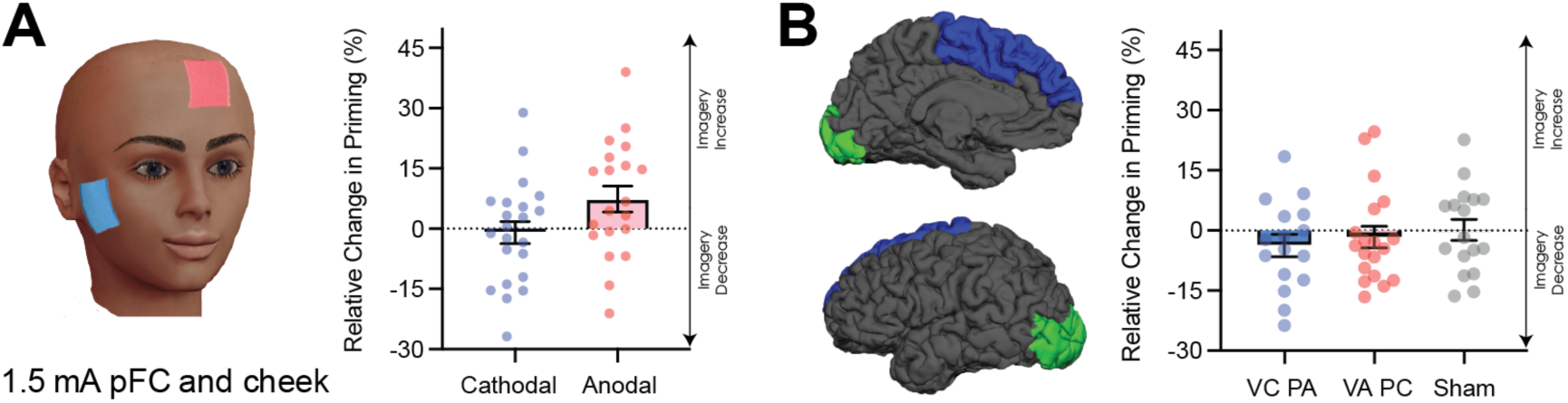
Data for prefrontal cortex stimulation. **A.** Effect of left prefrontal cortex stimulation on imagery strength at 1.5mA. The left image shows the tDCS montage, with the active electrode between Fz and F3 and the reference electrode on the right cheek. The right image shows the effect of cathodal (decrease excitability, blue dots represent each participant’s difference score) and anodal (increase excitability, red dots represent each individual participant’s difference score) stimulation averaged across all blocks during and after tDCS stimulation (D1, D2, P1 and P2). Imagery strength can be seen to increase with anodal stimulation. **B.** Effect of joint electrical stimulation of prefrontal cortex and visual cortex. Left image shows brain areas targeted in the final tDCS study. Data shows effect of cathodal occipital + anodal pFC stimulation (blue bars, blue dots represent individual participants data), anodal occipital + cathodal pFC stimulation (red bars, red dots represents individual participants data) and sham stimulation (grey bars, and grey dots represent individual participants data). All error bars show ±SEMs.

### Manipulating prefrontal cortex excitability

To explore the theoretical role of frontal cortex in imagery generation and maintenance further, we next sought to evaluate the effect of modulating neural excitability in prefrontal cortex using tDCS during image generation. The active electrode was placed between F3 and Fz (left frontal cortex), and the reference electrode on the right cheek (Fig.3A for montage). Participants completed both cathodal and anodal conditions (1.5mA) over two separate days. Interestingly, in contrast to the visual cortex where decreasing excitability led to stronger imagery, we found the opposite pattern for frontal areas. A linear mixed effects analysis was run with a 2 (tDCS polarity: cathodal and anodal), x 4 (block:D1, D2, P1, P2 – see S4A for timeline and S4G for data for each block) x 2 (order of stimulation: cathodal on first or second day) design. The effect of tDCS polarity was significant (χ^2^(1) = 6.1978, *p* = .01279, see figure 3A), with anodal

### The joint role of visual and frontal cortex activity in visual imagery strength

Beyond the individual roles of prefrontal and visual cortex in forming mental images, evidence suggests that both areas can act together as part of an imagery network (*20, 21*). Hence, we combined the whole-brain normalized mean fMRI intensity scores from the two areas (frontal and visual) and related their ratio to imagery strength. We found that the ratio of V1 to superior frontal predicted the strength of visual imagery (Spearman rank: r_s_ = -.53, *p* = .002). This effect also held when controlling for the Euklidean distance between the two areas (partial Spearman rank: r_s_ = -.54, *p* = .002). Hence, participants with both comparatively lower levels of visual cortex normalized mean intensity and higher frontal levels had stronger imagery.

To assess the possibility that cortical connectivity might be driving this fronto-occipital excitability relationship, we analyzed the individual functional connectivity of the same two areas for each participant, that is, the degree to which the BOLD signals in each area correlate over time. The functional connectivity did not significantly predict imagery strength (r = -.24, p = .19). This suggests that the combination of highly active frontal areas and low visual cortex excitability might present an optimal precondition for strong imagery creation, irrespective of the temporal coupling of their activity.

To further investigate this possibility, we ran a new tDCS experiment where both pre-frontal and visual cortex were simultaneously stimulated during imagery generation using the same blocked design as in all previous tDCS experiments (1.5mA). There were 3 conditions in this study, the first condition increased pre-frontal (anodal) and decreased visual cortex (cathodal) excitability, the second condition decreased pre-frontal (cathodal) and increased visual cortex (anodal) excitability, and the third condition was a sham condition where the tDCS machine shut off after 30 seconds. A linear mixed effects analysis was run with a 3 (tDCS polarity: cathodal, anodal and sham), x 4 (block:D1, D2, P1, P2 – see S5A for timeline and S5E for data for each block) x 3 (order of stimulation: cathodal on first, second or third day) design. The effect of tDCS polarity was not significant (tDCS χ^2^(2) = .36, *p* = .84)

In summary visual cortex excitability reliably correlated negatively with the strength of visual imagery using both fMRI and TMS as measurement tools (figure 1D-H). Modulating visual cortex excitability (specifically decreasing visual cortex excitability lead to increased visual imagery strength) also altered the strength of visual imagery (figure 2C & D, see table 1 for summary of all tDCS experiments). There was also evidence that altering pre-frontal cortex excitability modulates visual imagery strength, but in the opposite pattern to visual cortex – increasing pre-frontal cortex excitability led to increased imagery strength. However combining stimulation of the frontal and occipital cortex had no effect on modulating visual imagery strength.

**Table 1:** Summary of montage, intensity, duration and significance of each tDCS experiment.

| Experiment # | Montage (EEG Coordinates) | Intensity + Duration | Notes | Significant |
| --- | --- | --- | --- | --- |
| 1<br>Occipital | Active: Inion (Oz)<br>Reference: Supraorbital (Fpz) | 1 mA<br>15 minutes | Tested effect on imagery<br>strength | ✗ |
| 2<br>Occipital | Active: Inion (Oz)<br>Reference: Right Cheek | 1.5 mA<br>15 minutes | Tested effect on imagery<br>strength | ✓ |
| 3<br>Occipital | Active: Inion (Oz)<br>Reference: Right Cheek | 1.5 mA<br>15 minutes | Tested effect on imagery<br>strength (additional sham<br>condition) | ✓ |
| 4<br>Prefrontal | Active: Between F3-Fz<br>Reference: Right Cheek | 1.5 mA<br>15 minutes | Tested effect on imagery<br>strength | ✓ |
| 5<br>Occipital +<br>Prefrontal | Active: Inion (Oz)<br>Active: Between F3-Fz | 1.5 mA<br>15 minutes | Tested effect on imagery<br>strength | ✗ |
| Additional control<br>Occipital | Active: Inion (Oz)<br>Reference: Right Cheek | 1.5 mA<br>15 minutes | Tested effect on<br>phosphene threshold | ✓ |

## Discussion

Perhaps as far back as Plato, but overtly since the 1880s philosophers, scientists and the general populace have wondered why the human imagination differs so profoundly from one individual to the next. This question has recently gained fresh notability and attention with the introduction and classification of the term aphantasia to describe individuals who self-report no imagery at all (*22, 23*). Here we show the first evidence that pre-existing levels of neural excitability and spontaneous resting activity in visual cortex can influence the strength of mental representations. Our data indicate that participants with lower excitability in visual cortex have stronger imagery. Furthermore, we provide causative evidence, using tDCS over visual cortex, that altering neural excitability in these areas can modulate imagery strength. Prefrontal cortex excitability also played a role in controlling the strength of visual imagery, but in the opposite direction to visual cortex excitability.

The findings in visual cortex could be explained by hyperexcitability acting as a source of noise, which, when reduced, allows for a higher signal-to-noise ratio in visual cortex and thus stronger imagery. This hypothesis is in line with findings from related research. A study on grapheme-color synesthesia found that contradictory to our results synesthetes had enhanced resting state visual cortex excitability (measured using phosphene thresholds). However, they also found that synesthetic experience could be enhanced by reducing visual excitability via tDCS (*24*). These seemingly contradictory results were thought to be due to two different mechanisms. The authors suggested that a hyperexcitable visual cortex during brain development may be what leads to an individual developing synesthesia in the first place; in adulthood, however, decreasing visual cortex excitability might lead to increased signal-to-noise in the visual cortex, thereby enhancing the synethetic experience (*24*). In addition to the synesthesia research, the expectation of a visual stimulus leads to an imagery-like stimulus template, with reduced activity in V1 and improved stimulus decoding by pattern classifiers (*25*). Similarly, reduced early visual cortex activity increases the likelihood of visual hallucinations in a subsequent detection task (*26*). Further, behavioral data suggests that the presence of uniform afferent visual stimulation during mental image generation and visual working memory storage (*9, 10*) attenuates sensory strength and retention (*6, 9, 10*). The convergence of these data appear to indicate that ‘background’ neural noise in sensory cortices may play an important role in modulating the strength of mental representations.

Despite much evidence for the involvement of the prefrontal cortex and visual cortex working in concert during visual imagery we found that while increasing pre-frontal excitability by itself lead to increases in imagery strength, combining stimulation of visual and prefrontal cortex had no effect on visual imagery. It may be that in this condition an additional region, other than visual and pre-frontal cortex, was being modulated by our stimulation montage, leading to a null effect, or that this montage lead to smaller current densities and changes in excitability in both visual and pre-frontal cortex. Another possible explanation for these results is that modulating activity in two regions of the brain, is too much of a change and disrupts imagery formation, however neither of the stimulation conditions resulted in significant reductions as compared to sham stimulation making this explanation unlikely. There also exists large variability in pre-frontal cortex anatomy and tDCS effectiveness. Recent research has shown that the thickness of left prefrontal cortex correlated with behavioural changes from anodal (but not cathodal) stimulation(*27*). It might be that these large variations might play a role in our null findings, however we did find that isolated stimulation to pre-frontal cortex modulated imagery strength, making this also an unlikely explanation of these null results.

Our findings do conflict with some previous research on visual imagery. For example one study found that applying 1Hz of TMS to area BA 17 (primary visual cortex), slowed responses on a task where individuals had to imagine some stripes (or were perceptually shown stripes) and answer questions about these images(*28*). Although these chronometry type experiments are very common in early visual imagery research, and were important in advancing the field as a whole, they do not provide any information about the quality or sensory representational nature of the images held in mind. Slower reaction times on both the perception and imagery task may be due to a general slowing of cognitive performance or visual scanning, rather than reflecting any change in the quality of the visual images created in the mind. Previous work has also found positive correlations between BOLD activity in the visual cortex and the vividness of visual imagery questionnaire (*29, 30*). Additionally some TMS studies have found that during visual imagery, visual cortex excitability increases(*31*). These findings at first may seem contradictory however these tasks are all online tasks, and to calculate these BOLD changes a baseline of ‘resting-state’ activity is used. It may be that participants with initially low visual cortex excitability are able to increase visual cortex activity more-so than those with higher-levels, and this could potentially explain the larger BOLD changes for individuals with stronger visual imagery.

It is possible that the observed effects of cortical excitability may be driven by individual differences in inhibitory and excitatory neurotransmitter concentrations. Numerous studies have investigated what neurotransmitters modulate cortical excitability with GABA and Glutamate being implicated in controlling inhibition and excitability, respectively. The concentration of glutamate in the early visual cortex has shown to correlate positively with visual cortex excitability (measured by phosphene thresholds) in both normal and synaesthetic participants; in contrast, no relationship between GABA and cortical excitability was found(*32*). In addition, there is evidence for a strong link between BOLD-fMRI activity and glutamate concentration: using a combined fMRI-MRS approach where BOLD-fMRI activity and glutamate signals were recorded simultaneously, researchers found that the time courses of fMRI-BOLD activity and Glutamate concentration were strongly correlated (*33*). Evidence whether such a relationship also holds on a between-subject level is still missing, but seems plausible. If this is the case, then the observed relationships of our neural measures and visual imagery may (at least partly) be due to individual differences in the concentration of glutamate in visual cortex: a lower level of glutamate in the visual cortex might result in less excitatory neuronal noise, thereby increasing the signal-to-noise ratio of top-down signals that govern the generation of internal images in the visual cortex.

As found in our study, tDCS was able to manipulate visual imagery strength. It is possible this may be due to the up- or down-regulation of concentrations of glutamate. However, the exact mechanisms underlying the change in cortical excitability during brain stimulation is still not well understood. Previous studies found a myriad of changes in neurotransmitter releases, as well as changes in neuromodulator concentrations(*16, 34*). Additionally, the mechanisms of change in cortical excitability are thought to be different during and after stimulation(*34*): Altered cortical excitability *during* tDCS stimulation is thought to be due solely to changes in membrane potential(*34*), whereas the mechanisms of the changes *after* tDCS stimulation are thought to be very similar to long-term potentiation (increasing excitability) and long term depression (decrease excitability). These changes likely rely on the way anodal and cathodal stimulation differentially displace Magnesium ions on the NMDA ion channels resulting in either a rise (anodal) or decrease (cathodal) in post-synaptic Ca2+ concentrations (*34*).

It is important to note that while tDCS has been shown to modulate visual cortex excitability in numerous studies as well as our control experiment, there are large individual differences in the amount of change that occurs. This may be due to numerous individual factors, such as skull thickness, resting state excitability, and hormonal levels(*35, 36*). In our study, although the majority of participants showed the expected pattern of results (larger increases in imagery strength in the cathodal vs anodal condition for occipital stimulation), there were some participants who showed the opposite pattern of results. Why some participants benefit from tDCS and others do not was not in the scope of the current experiment; future experiments using online measurements of cortical excitability changes (e.g. EEG) will help to address this interesting and important question.

Over the last 30 years empirical work has demonstrated many commonalities between imagery and visual perception (see (*2, 3*) for a review). However, the two experiences have clear phenomenological differences between them. Our findings suggest a possible novel dissociation between mental imagery and visual perception, as perceptual sensitivity is associated with higher levels of visual cortex excitability, whereas our results suggest the opposite for mental imagery; stronger imagery is associated with lower visual excitability.

A plethora of imagery research has demonstrated evoked and content specific BOLD responses in early and later visual cortex when individuals form a mental image (*2, 3*). Here however, we took a different approach by examining the individual variation in brain physiology that might form the preconditions for strong or weak imagery. This endeavor required a non-event related design. Interestingly, such non-event related designs utilizing inter-individual differences are now commonly used to mechanistically link human cognition and brain function or anatomy (*37*). Our results add to this growing body of research, which demonstrates that pre-existing brain activity parameters can fundamentally influence mental performance.

Our data suggest that neural excitation in visual cortex, plays a key role in governing the strength of mental imagery. Our observations may also have clinical applications: In many mental disorders, imagery can become uncontrollable and traumatic. On the other hand, mental imagery can also be harnessed specifically to treat these disorders (*2*). Interestingly, disorders that involve visual hallucinations such as schizophrenia and Parkinson’s disease are both associated with stronger and/or more vivid mental imagery (*38, 39*). It has recently been suggested that the balance between top-down and bottom-up information processing may be a crucial factor in the development of psychosis, with psychosis prone individuals displaying a shift in information processing towards top-down influences over bottom-up sensory input (*40*). Our data indicate it may be possible to treat symptomatic visual mental content by reducing its strength via non-intrusively manipulating cortical excitability. Alternatively, we may be able to ‘surgically’ boost mental image simulations specifically during imagery-based treatments, resulting in better treatment outcomes. Further research on longer lasting stimulation protocols, and the individual differences in response to brain stimulation, is needed to assess its therapeutic potential.

In conclusion our data demonstrates that visual cortical excitability appears to play a role in governing the strength of an individual’s visual imagery strength providing a potential explanation for the large variation in visual imagery that exists within the general population and providing a potential new tool for altering the strength of visual imagery.

## Materials and Methods

### Study Design

The first study with fMRI was exploratory, to assess whether resting levels of BOLD might predict visual imagery. We followed this with a correlational TMS study and aimed to collect phosphene thresholds from 30-35 participants, which would give us power of around 80-85% for a moderate correlation (in line with the MRI correlations of r =∼ .45). All tDCS experiments were designed as repeated measures studies (with the aim of all participants completing all conditions in the study). We aimed to collect data from 15-20 participants, as most tDCS studies examining effects on cognition have found significant effects with this range of participants (See for examples: (*44–47*)). Data collection stopped once we had at least 15 participants in each group who had completed 2 days of testing, no more participants were recruited beyond this point, however participants were not cancelled if we reached 15 (e.g. if we had collected 15 participants for both days but still had 2 more participants who had completed one day of testing, we still ran them through the study – resulting in 17 participants).

### Participants

All MRI participants were right-handed and had normal or corrected-to-normal vision, with no history of psychiatric or neurological disorders. All tDCS and TMS participants had normal or corrected-to-normal vision, with no history of psychiatric or neurological disorders, as well as no history of migraines and/or severe or frequent headaches. All MRI research was carried out in Germany at the Max Planck Institute for Brain Research and all brain stimulation research (tDCS and TMS) was carried out in Australia at the University of New South Wales. Written informed consent was obtained from all participants and the ethics committee of the Max Planck Society approved the MRI study and the ethics committee of the University of New South Wales approved the tDCS and TMS studies.

### Exclusion Criteria

There were a number of strict exclusion criteria chosen a priori due to the technical psychophysics and brain stimulation experiments involved here. These are based on previous work using the binocular rivalry paradigm, which is a sensitive measure of visual imagery strength when participants complete the task correctly. Due to the nature of the task it is important to include catch trials/exclusion criteria to assess whether participants are correctly and reliably completing the task. These values are based on exclusion criteria we have used in previous experiments using this paradigm.

#### Brain imaging sample

32 individuals (age range: 18 - 36 years, median: 25.5; 13 males) participated in the fMRI resting-state and retinotopic mapping measurements and in the behavioral experiment. These individuals had been part of previous studies(*5, 8*). Of the original imagery study(*5*), which included 34 participants, 1 participant had not done the fMRI resting-state measurement (but this individual participated in the additional fMRI resting-state measurement, see further below). The other participant was excluded because of reporting several migraine attacks shortly prior to the measurement. Migraine is known to affect fMRI BOLD activity and cortical excitability (*42, 43*). One participant was excluded from data analysis because of misunderstanding the task instructions in the behavioral imagery task (this participant had already been excluded in the original study). Of the remaining 31 individuals, a subsample of 22 also participated in the luminance condition of the behavioral experiment, which was conducted in a separate session. To look at the reliability of the observed relationships, we also analysed the data of an additional fMRI resting-state measurement (with different sequence parameters, see further below). This sample included 31 individuals (age range: 22-36 years). 30 of these had also participated in the original resting-state measurement. After excluding the above-mentioned participant who had misunderstood the task instructions of the behavioural task, a surface-based group analysis on the brain-behaviour relationships was computed using the remaining 30 individuals. Participants were reimbursed for their time at a rate of 15€ per hour. Written informed consent was obtained from all participants and the ethics committee of the Max Planck Society approved the study.

#### TMS samples

All participants in both the TMS and tDCS studies had normal or corrected to normal vision, no history of any neurological or mental health issues or disorders, no history of epilepsy or seizures themselves or their immediate family, no history of migraines and no metal implants in the head or neck region. We aimed to collect phosphene thresholds from 30-35 participants, which would give us power of around 80-85% for a moderate correlation (in line with the MRI correlations). A total of thirty-seven participants participated in this study for money ($30 per hour) or course credit, five participants were excluded due to an inability to produce reliable phosphenes (see table 2 and exclusion criteria). Of the remaining thirty-two participants 15 were female, age range: 18-30). Written informed consent was obtained from all participants and the ethics committee of the University of New South Wales approved the study.

**Table 2:**
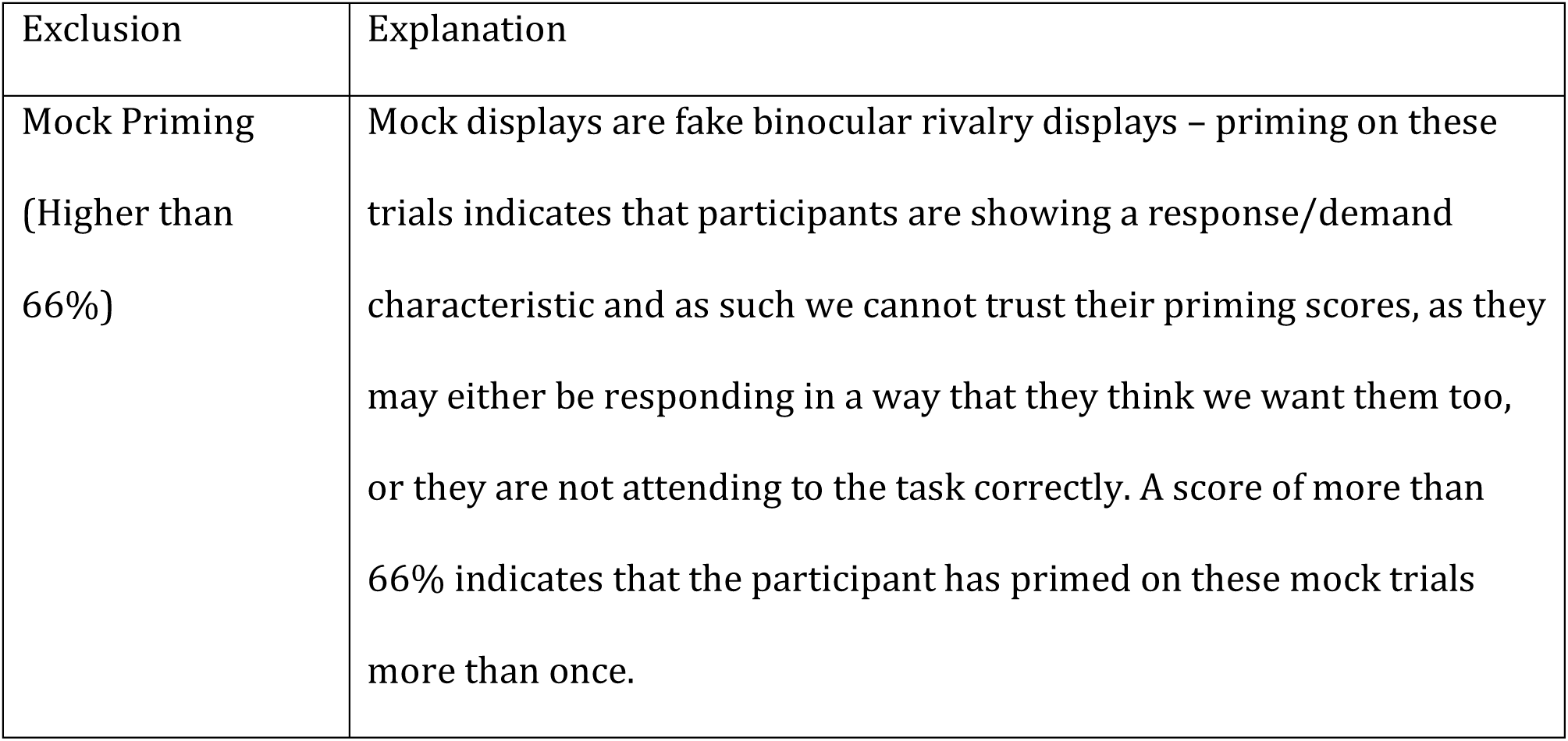
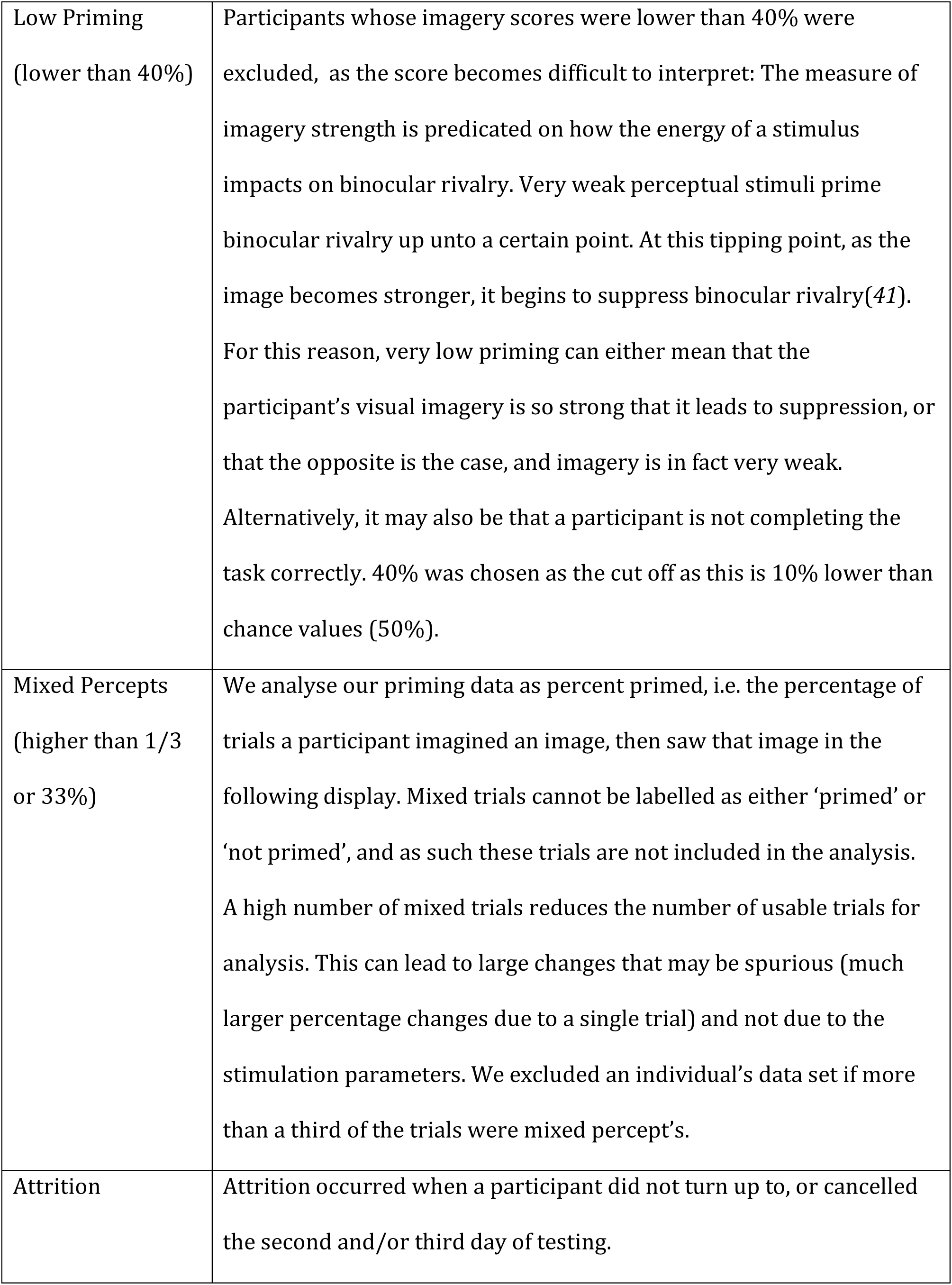
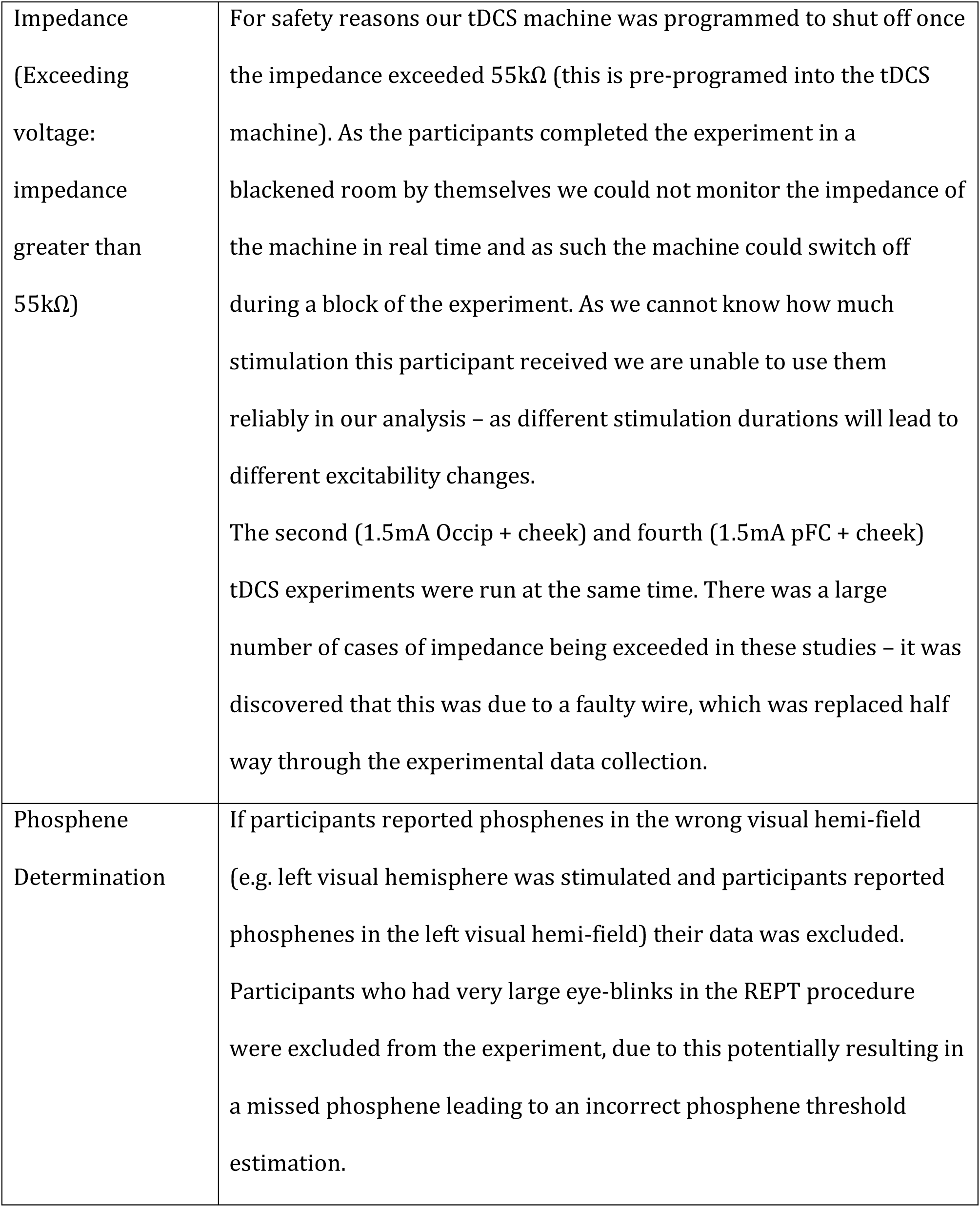
Exclusion criteria

#### TDCS samples

For all tDCS experiments we aimed to collect data from 15-20 participants, as most tDCS studies examining effects on cognition have found significant effects with this range of participants (See for examples: (*44–47*)). A priori we chose a cut-off of 33% of trials being mixed as an exclusion criteria (see table 1 for explanations of all exclusion criteria). The reason for this is that mixed trials are not included in our analysis, and as such a large number of mixed trials vastly decreases the number of analyzable trials. Participants whose imagery scores were lower than 40% were also excluded, as it is difficult to tell if these data should be defined as strong or weak imagery, or due to a participant just not completing the task correctly. This measure of imagery strength is predicated on how the energy of a stimulus impacts on binocular rivalry. Very weak perceptual stimuli prime binocular rivalry up unto a certain point. At this tipping point, as the image becomes stronger, it begins to suppress binocular rivalry(*41*). For this reason when priming is low this indicates suppression that may either mean the participant’s visual imagery is so strong it is suppressing binocular rivalry, or the participant is not completing the task correctly.

For the first tDCS experiment (1mA, Occipital and Supraorbital) a total of twenty-one subjects participated for money or course credit. Five participants were excluded from our analysis, as the number of usable trials was small due to too many reported mixed rivalry percepts (more than a third of trials, N = 3) and two were excluded due to low priming scores (see table 1 for exclusion criteria). Of the remaining sixteen participants 7 were female, and the age range was 18-32.

For the second tDCS experiment (1.5mA, Occipital and Cheek) a total of thirty-nine subjects participated for course credit. Due to a faulty tDCS cable many of these participant’s had the tDCS machine shut off/exceeded voltage on the first day of testing (N = 9, see table 2 for exclusion criteria) and they did not come back for a second day of testing. Of the 30 participants who did not exceed voltage in the first day two were excluded due to low priming and two had a high number of mixed trials (see table 1 for exclusion criteria). Of the remaining 26, eight participants only had one day of testing data available due to the machine exceeding voltage on the second day of testing, two participants had one day of testing removed due to low priming on one of the two days, age range 18-26.

For the third tDCS experiment (1.5mA, Occipital and Cheek + Sham) a total of 28 subjects participated for course credit or payment ($40 per hour). Two participants were excluded due to technical issues with the computer on the first day of testing, one participant was excluded for pressing the incorrect buttons during the task/due to misunderstanding of the task, one participant was excluded due to removing the tDCS during the testing session, one participant was removed for a high number of mixed trials and one for high mock priming (see table 1 for exclusion criteria). Of the remaining 22 participants age range was 18-23. Of these participants 4 only completed 2 of the 3 days of testing due to attrition (N = 3) and machine malfunction/exceeded voltage (N = 1). Another 4 participants also only completed one day of testing due to attrition (N = 3) and machine malfunction (N = 1).

For the fourth tDCS experiment (1.5mA, left prefrontal and cheek) thirty-one participants participated in the study for course credit. Due to a faulty tDCS cable many of these participant’s the tDCS machine shut off/exceeded voltage on the first day of testing (N = 4), and as such they did not complete the study. Two participants’ data was removed due to very high mock priming, indicating either a misunderstanding of the task or demand characteristics and one was removed due to very low priming (see table 1 for exclusion criteria). Of the remaining twenty four participants seven had one day’s worth of data due to the tDCS machine shutting off/exceeding voltage (N = 3) or attrition (N = 2), or having very low imagery priming on one of the days (N = 2), age range was 18-25 years.

For the fifth tDCS experiment (1.5mA, left prefrontal and occipital cortex + sham) twenty eight participants participated in the study for course credit or payment (AUD $40 per hour). Three participants were excluded due to very low priming, one participants was excluded due to technical issues with the tDCS machine (exceeding impedance/voltage on the first day) and five participants were excluded due to misunderstandings or incorrectly completing the task (either pressing incorrect buttons (N = 3) or 100% priming for mock rivalry (N =2), see table 1 for exclusion criteria). Of the remaining nineteen participants the age range was 18-35 years and of these participants one participant did not complete the sham condition (machine malfunction – exceeded impedance/voltage) and three participants did not complete the occipital cathodal + prefrontal anodal condition (machine malfunction– exceeded impedance/voltage).

All subjects participated in these studies for course credit or money - $30 AUD per hour.

#### tDCS modulation of phosphene thresholds control study

A total of twenty-nine subjects participated in this study for money ($30 AUD per hour) or course credit. Of these 29 participants eleven were excluded due to a number of strict exclusion criteria in regards to reliability of phosphene thresholds: If participants reported phosphenes in the wrong visual hemi-field (e.g. left visual hemisphere was stimulated and participants reported phosphenes in the left visual hemi-field) their data was excluded (N = 2), see table 1 for exclusion criteria. If participants blinked during the rapid estimation of phosphene thresholds (REPT) procedure their data was also removed from analysis (N = 4). A participant’s data was also removed if the REPT procedure took longer than five minutes to set up after tDCS stimulation (N = 1). 3 participants were also removed due to technical issues with the tDCS machine exceeding voltages on one of the days (N = 2) or the REPT Matlab procedure experiencing errors (N = 1). One participant was removed due to attrition (N = 1). This resulted in 18 participants’ data being analysed (8 female, age range 18-25).

#### TMS Phosphene threshold reliability study

This sample consisted of the same twenty-nine subjects that participated in the control study to test tDCS modulation of phosphene thresholds. Exclusion criteria were the same as stated above, with the exception that those participants who had technical issues with the tDCS machine were included as their pre-tDCS TMS phosphene values were still usable (N=2); the study also included those 1 participant who took longer than 5 minutes to set up the TMS after the tDCS stimulation. This resulted in 21 participants’ data being included in this correlation.

## Behavioral measurements

### Apparatus

#### Brain imaging sample

Participants sat in a darkened room with dark walls, wearing red-green anaglyph glasses for the binocular rivalry imagery paradigm. Their head position was stabilized with a chin rest and the distance to the screen was 75 cm. The stimuli were presented on a CRT monitor (HP p1230; resolution, 1024 x 768 pixels, refresh rate: 150 Hz; visible screen size: 30° x 22.9°) and controlled by MATLAB R2010a (The MathWorks, Natick, MA) using the Psychophysics Toolbox extension (*48–50*), running on Mac OSX, version 10.7.4.

#### TMS/tDCS sample

All experiments were performed in a blackened room on a 27 inch iMac with a resolution of 2560×1440 pixels, with a frame rate of 60Hz. A chin rest was used to maintain a fixed viewing distance of 57cm. Participants wore red-green anaglyph glasses throughout all experiments.

### Stimuli

#### Brain imaging sample

The circular Gaussian-windowed Gabor stimuli were presented centrally, spanning a radius of 4.6° around the fixation point in visual angle (thereby covering a diameter of 9.2°), one period subtending a length of 1.2°. The peak luminance starting value was ∼0.71 cd/m^2^ for the red horizontal grating, and ∼0.73 cd/m^2^ for the green vertical grating, which was then individually adjusted for each participant to compensate for eye dominance (see further below).

#### TMS/tDCS sample

The binocular rivalry stimuli were presented in a Gaussian-windowed annulus around the bull’s eye and consisted of a red-horizontal (CIE X = .579 Y = .369 and green-vertical (CIE X = .269 Y = .640), Gabor patch, 1 cycle/°, with a diameter of 6° and a mean luminance of 6.06cd/m^2.^ The background was black throughout the entire experiment.

#### Mock trials

Mock rivalry displays were presented on 10% of trials in the behavioral measurements with the brain imaging sample, 25% of trials in the first two tDCS experiments as well as the TMS experiment, and in 12.5% of the third tDCS experiment to assess demand characteristics. The mock displays consisted of a spatial mix of a red-horizontal and green-vertical Gabor patch (50/50% or 25/75%). The mock display was spatially split with a blurred edge and the exact path of the spatial border changed on each trial based on a random function. Otherwise the mock rivalry displays had the same parameters as the Gabor patches described in the previous paragraph.

### Procedure

All participants first underwent a previously documented eye dominance task(*6*). They then underwent the binocular rivalry imagery paradigm which has been shown to reliably measure the sensory strength of mental imagery through its impact on subsequent binocular rivalry perception (*1, 6, 7, 9, 10, 51-53*), thus avoiding any reliance on self-report questionnaires or compound multi-feature tasks. Previous work has demonstrated that when individuals imagine a pattern or are shown a weak perceptual version of a pattern, they are more likely to see that pattern in a subsequent brief binocular rivalry display (see (*1*) for a review). Longer periods of imagery generation or weak perceptual presentation increase the probability of perceptual priming of subsequent rivalry (*6, 53, 54*). For this reason, the degree of imagery priming has been taken as a measure of the sensory strength of mental imagery(*1, 6, 7, 9, 10, 51-53*). This measure of imagery strength has been shown to be both retinotopic location- and spatial orientation-specific (*15, 51*), is closely related to phenomenal vividness (*7, 55*), is reliable when assessed over days (*7*) or weeks (*15*), is contingent on the imagery generation period (therefore not due to any rivalry control (*52*)) and can be dissociated from visual attention (*6*).

At the beginning of each trial of the imagery experiment, participants were presented with a letter ‘R’ or ‘G’ which indicated which image they were to imagine (R = red-horizontal Gabor patch, G = green-vertical Gabor patch). Participants then imagined the red or green pattern for either six (tDCS and TMS experiments) or seven seconds (behavioral measurements of the brain imaging sample). Following this imagery period, the binocular rivalry display appeared for 750ms and participants indicated which image was dominant by pressing ‘1’ for mostly green, ‘2’ for a mix and ‘3’ for mostly red. During the behavioral measurements of the brain imaging sample and in both the 1.5 ma occipital tDCS experiments, as well as the 1.5ma pFC tdcs experiment, on-line ratings of imagery vividness were collected by having participants rate the vividness of the image they had created (on a scale of ‘1’ = least vivid to ‘4’ = most vivid) on each trial after the imagery period and before the binocular rivalry display.

For the tDCS experiments there were no effects for mean subjective ratings of imagery vividness (see Supplementary Fig. S1C). For the subjective vividness ratings acquired in the brain imaging sample, we conducted a whole-brain surface-based analysis of the fMRI resting-state data (see Methods and Supplementary Fig.S1B and Table S3). During the imagery experiment where background luminance was included, the procedure was the same as the basic imagery experiment, except that during the imagery period, the background ramped up and down (1s up, 1s down, to avoid visual transients) to a yellow color (the mean of the two binocular rivalry colors). Throughout all imagery experiments, participants were asked to maintain fixation on a bulls-eye fixation point in the center of the screen.

#### Brain imaging sample

Participants completed 100 trials of the standard imagery paradigm per session (outside the scanner). The behavioral test session was repeated after an average of ∼2 weeks with each participant. All of the runs were divided into blocks of 33 trials, and participants were asked to take a rest in between. In one participant, there was a strong perceptual bias for 1 of the 2 rivalry patterns in the first session due to incorrectly conducted eye dominance adjustments. Therefore, only the data set from the second session of this participant was used for later analysis. The retests demonstrated a very high retest reliability of the imagery strength measure (*r* = .877, *p* < .001). In the luminance condition, which was tested on a subsample of the original sample in another session, the participants completed 50 trials. The data from both conditions were checked for normal distribution using Shapiro-Wilk normality test. No violation of the normality assumption was detected (both *p* > .52).

#### TMS/tDCS samples

For the TMS study, participants completed one block of 40 imagery trials. In the first tDCS experiments comprising of cathodal and anodal conditions, participants completed a total of 40 trials for each block resulting in a total of 480 trials across the two days of testing. In the tDCS experiments with cathodal, anodal and sham conditions participants completed 48 trials per block, and as such 864 trials were completed.

#### Control tDCS modulation of phosphene thresholds experiment

Participants completed both the anodal and cathodal stimulation across two days separated by at least 24 hours, the order of which was randomized and counterbalanced across participants. Participants completed a memory or psychophysical task (both of which are not relevant to the current study) followed by the automated REPT phosphene threshold procedure prior to tDCS stimulation. Following this, participants completed two blocks of the imagery task (see main texts methods for full description of procedure and stimuli) with fifteen minutes of cathodal or anodal stimulation. Immediately after the tDCS stimulation participants completed the automated REPT procedure again.

### Neuroimaging experiments

All neuroimaging data were acquired at the Brain Imaging Center Frankfurt am Main, Germany. The scanner used was a Siemens 3-Tesla Trio (Siemens, Erlangen, Germany) with an 8-channel head coil and a maximum gradient strength of 40 mT/m. Imaging data were acquired in two or three scan sessions per participant..

#### Anatomical imaging

For anatomical localization and coregistration of the functional data, T1-weighted anatomical images were acquired first using an MP-RAGE sequence with the following parameters: TR = 2250 ms, TE = 2.6 ms, flip angle: 9°, FoV: 256 mm, resolution = 1 x 1 x 1 mm^3^.

### fMRI Retinotopic mapping measurement and analysis

This procedure has already been described in previous studies (*15, 56, 57*); the retinotopic maps acquired in studies (*^5, 8^*) were used for the ROI-based fMRI resting-state analyses in the present study. A gradient-recalled echo-planar (EPI) sequence with the following parameter settings was applied: 33 slices, TR = 2000 ms, TE = 30 ms, flip angle = 90°, FoV = 192 mm, slice thickness = 3 mm, gap thickness = 0.3 mm, resolution = 3 x 3 x 3 mm^3^. A MR-compatible goggle system with two organic light-emitting-diode displays was used for presentation of the stimuli (MR Vision 2000; Resonance Technology Northridge, CA), which were generated with a custom-made program based on the Microsoft DirectX library (*58*). The maximal visual field subtended 24° vertically and 30° horizontally.

#### Retinotopic mapping procedure

To map early visual cortices V1, V2 and V3, our participants completed two runs, a polar angle mapping and an eccentricity mapping run. The rationale of this approach has already been described elsewhere (*59, 60*). Polar angle mapping: For the mapping of boundaries between areas, participants were presented with a black and white checkerboard wedge (22.5° wide, extending 15° in the periphery) that slowly rotated clockwise around the fixation point in front of a grey background. In cycles of 64 s, it circled around the fixation point 12 times at a speed of 11.25 in polar angle/volume (2 s). Eccentricity mapping: To map bands of eccentricity on the cortical surface to the corresponding visual angles from the center of gaze, our participants were presented with a slowly expanding flickering black and white checkerboard ring in front of a grey background (flicker rate: 4 Hz). The ring started with a radius of 1° and increased linearly up to a radius of 15°. The expansion cycle was repeated 7 times, each cycle lasting 64 s. The participants’ task in both mapping experiments was to maintain central fixation.

#### Retinotopic mapping data analysis

We used FreeSurfer’s surface-based methods for cortical surface reconstruction from the T1-weighted image of each participant (*61, 62*) (http://surfer.nmr.mgh.harvard.edu/fswiki/RecommendedReconstruction). FSFAST was applied for slice time correction, motion correction and co-registration of the functional data to the T1-weighted anatomical image. Data from the polar angle and eccentricity mapping experiment were analysed by applying a Fourier transform to each voxel’s fMRI time series to extract amplitude and phase at stimulation frequency. Color-encoded F-statistic maps were then computed, each color representing a response phase whose intensity is an F-ratio of the squared amplitude of the response at stimulus frequency divided by the averaged squared amplitudes at all other frequencies (with the exception of higher harmonics of the stimulus frequency and low frequency signals). The maps were then displayed on the cortical surface of the T1-weighted image. Boundaries of areas V1, V2 and V3 were then estimated manually for each participant on the phase-encoded retinotopic maps up to an eccentricity of 7.2°.

### fMRI Resting-state data acquisition and analysis

#### fMRI resting-state data acquisition

The fMRI resting-state data (TR2) were collected using a gradient-re-called echo-planar imaging (EPI) sequence with the following parameters: TR = 2000 ms, TE = 30 ms, flip angle = 90°, FoV = 192 mm, slice thickness = 3 mm, number of slices = 33, gap thickness = 0.3 mm, voxel size = 3 x 3 x 3 mm^3^, acquisition time = 9 minutes, 20 seconds (thus, 280 volumes were collected)The additional fMRI measurement (TR1) used a gradient-recalled echo-planar imaging (EPI) sequence with the following parameters: TR = 1000 ms, TE = 30 ms, flip angle = 60°, FoV=210mm, slice thickness = 5mm, number of slices = 15, gap thickness = 1 mm, voxel size = 3.28 x 3.28 x 5 mm^3^, acquisition time = 7 minutes, 30 seconds (i.e. 450 volumes). During the scans, the screen remained grey and participants had no further instruction but to keep their eyes open and fixate a cross in the center of the grey screen. Of those individuals who participated in both fMRI measurements, half completed the two resting-state measurements in the same session, whereas the other half completed the measurements on two different days.

#### fMRI resting-state data analysis: whole-brain surface-based group analysis

For a first assessment of the relationship between behavior and the fMRI data, we ran whole-brain analyses with the mean fMRI intensity data using a surface-based group analysis in FreeSurfer. Preprocessing of the functional data was done using FSFAST, which included slice time correction, motion correction and co-registration to the T1-weighted anatomical image. No smoothing was applied, and the first 2 volumes (TR2 data) or 4 volumes (TR1 data) of the fMRI measurement were discarded. In a first-level analysis, each individual’s average signal intensity maps were computed (which included intensity normalization) and nonlinearly resampled to a common group surface space (fsaverage), which allows for comparisons at homologous points within the brain. Following this, all subjects’ data were concatenated and a general linear model fit to explain the individual behavioral data by the individual mean fMRI intensity levels was computed vertex-wise using an uncorrected threshold of *P* < 0.05. Correction for multiple comparisons was done using a pre-cached Monte Carlo Null-Z simulation with 10 000 iterations and a cluster-wise probability threshold of *P* < 0.05.In addition, as we had also collected subjective vividness ratings in the brain imaging sample (see Procedure), we also ran the equivalent whole brain analysis for the vividness ratings with the TR2 data set, using each individual’s mean vividness (see Supplementary Fig .S1B and Supplementary Table S3). As already described in our previous study (*15*), the subjective vividness values of two individuals were extreme, leading to a violation of the normal distribution assumption. As normality is necessary for the general linear model fit (Shapiro-Wilk normality test: W(31) = .885, *p* = .003), the vividness ratings of these two individuals were excluded in the whole brain analysis.

#### fMRI resting-state data analysis: ROI-based approach

The fMRI resting-state data were first preprocessed individually for each participant using the preprocessing steps implemented in FSL’s MELODIC Version 3.10 (http://fsl.fmrib.ox.ac.uk/fsl/fslwiki/MELODIC), which included motion and slice time correction, high-pass temporal filtering with a cut-off point at 200 seconds and linear registration to the individual’s T1 anatomical image and to MNI 152 standard space. No spatial smoothing was applied. The first two of the 280 volumes of the TR2 measurement were discarded to allow for longitudinal magnetization stabilization (of the TR1 measurement, the first 4 volumes of the 450 volumes were discarded); further ROI-based analyses confirmed that the pattern of significant results in visual cortex did not change when more volumes (4, 6 and 8) were discarded (see Supplementary Figure S8). To compute fMRI mean intensity of the early visual cortex in each individual’s subject space, delineations of the areas were first converted from anatomical to functional space in each individual. To ensure that the conversion had not produced overlaps between areas V1-V3, the volumes were subsequently subtracted from each other. Time courses of V1-V3 were then determined to compute their mean intensity across time. To determine mean intensity for other brain areas, we relied on the gyral-based Desikan–Killiany Atlas (*63*). To ensure that there was no overlap between posterior atlas-defined areas and the retinotopically mapped early visual cortex, which would result in the mean intensity of these areas being partly computed from the same voxels, the volumes of the retinotopically mapped areas were also subtracted from the adjacent atlas-defined areas; the fMRI mean intensity of the atlas-defined areas was then determined from the remainder of these. The estimates of fMRI mean intensity of the atlas- and retinotopically mapped areas (*12*) were normalized by subtracting the whole brain’s mean intensity from the area’s mean intensity, divided by the standard deviation of the whole brain’s mean intensity.

Like the behavioral data, the normalized mean intensity values were checked for normal distribution using Shapiro-Wilk normality test. None of the retinotopically mapped early visual cortices showed a violation of the normal distribution (all *p* > .20). Of the 34 atlas-defined areas, the normalized mean fMRI intensities of 4 areas showed a violation of the normal distribution assumption (*p* < .05; fusiform, inferiortemporal, parstriangularis and postcentral area). For this reason, the relationships with behavior were also computed using Spearman rank correlations. Like with Pearson product moment correlations, none of the intensities of these areas had a significant relationship with behaviour (all *p* >.20). The ratio of V1 and superior frontal mean intensities showed a violation of the normal distribution assumption (W(31) = .919, *p* = .022) due to one extreme value (subject S8). Therefore, Spearman rank correlation (*r_s_*) was used to compute the relationship with behavior. To further examine the possibility that temporal coupling between V1 and superior frontal cortex might account for their inverse relationship with behavior, we also computed each individual’s functional connectivity of these two regions by calculating the time-wise correlation of their resting-state signals in each individual. As the functional connectivity data did not violate the normal distribution assumption (Shapiro-Wilk normality test, *p* = .497), Pearson product moment correlation was used to examine the relationship with behavior. To correct for multiple comparisons in the brain-behaviour relationships, p-values for adjusted using the False Discovery Rate (FDR)(*64, 65*). In the analysis where we used Steiger’s Z to compare brain-behaviour correlations of the standard imagery task to the imagery task with background luminance, permutation-based multiple comparison correction was done as follows: we first randomly shuffled the labels of the two behavioural tasks and re-computed the correlations and Fisher Z-transformed correlations with each ROI (1000 permutations). We then re-computed Steiger’s Z from all permutations, summarized the distributions from all ROIs and identified the levels of significance of the actual Steiger’s Z values by their percentile in the distribution.

#### Control analyses to examine the effect of individual head motion

To rule out the possibility that factors like individual differences in head motion contributed to intensity variability, we re-analysed the data in 2 ways: (1) Partialling out overall head movement: we first computed a measure of Framewise Displacement (FD), which is a frame-wise scalar quantity of the absolute head motion from one volume to the next using the 6 motion parameters (*66*). To obtain one value of absolute movement parameter for each individual, we determined the absolute sum of the displacement values across the whole run. We then used this parameter as a control variable in a Spearman partial correlation analysis (since the normality assumption was violated for overall head motion, *W*(31) =.790, *p* < .001). Exclusion of volumes with strong head movements (“scrubbing”): volumes with strong head movements have previously been shown to influence functional resting-state connectivity patterns (*66*). To investigate whether such movement also influenced individual mean resting-state intensity in our sample, we re-analysed our data as using the method described by Power et al. To identify affected timepoints (i.e. volumes), we first computed the Framewise Displacement (FD), and DVARS, which is a frame-wise measure of how much the signal changes from one volume to the next. Using Powell et al.’s threshold of 0.5 for FD and 0.5% signal change for DVARS, we then excluded all volumes that had FD and DVARS values above threshold, plus one frame back and two forward. After this procedure, we re-computed mean intensity for each participant, and correlated the values with behaviour.

### Phosphene Threshold Determination

Phosphene thresholds were obtained using single pulse TMS with a butterfly shaped coil (Magstim Rapid^&^, Carmarthenshire, UK). The coil was placed centrally and approximately 2 cm above the inion. To obtain each participant’s phosphene threshold, we used the previously documented automated rapid estimation of phosphene thresholds (REPT) (*67*). This REPT procedure uses a Bayesian adaptive staircase approach to find the 60% phosphene threshold of each participant.

Before the REPT procedure we first ensured that our participants were able to see reliable phosphenes. We initially placed the coil centrally and approximately 2 cm above the inion (with the coil oriented 45 degrees) and gave the participant’s single pulses starting at 50% of the machines maximal output, moving up to 85% of the machines maximal output in 5% step increments. If the participant failed to report any phosphenes (or if the location of the coil was producing very large eye blinks or neck twitches) the coil was moved approximately 1cm left and the same procedure occurred. If this did not elicit phosphenes the coil was moved to the right hand side (approximately 2cm above the inion and 1cm to the right) and the same procedure occured. If the participant still could not see any phosphenes they did not complete any more of the experiment. While this initial testing took place participants were seated in front of a computer screen with a piece of black cardboard with white numbered quadrants covering the monitor. The participants were instructed to relax and stare forward at a fixation dot in the middle of the black cardboard and to let the experimenter know if they saw any sort of visual disturbances on the cardboard and if they did see something to describe what they saw and in which quadrant they saw it. If participants reported a phosphene occurring in an incorrect location, e.g. the left visual field when stimulating the left visual cortex, then they were excluded from the study and did not complete the REPT procedure.

During the REPT procedure participants were seated in front of the same computer screen as the initial phosphene testing with a piece of black cardboard with white numbered quadrants covering the monitor. The coil was placed in the same location as where phosphenes were elicited in the initial phosphene testing described above using a clamp attached to the testing table. Participants received 30 pulses, of varying intensities, which were delivered automatically by the machine when the participant pressed the space key (self paced). After each pulse participants were instructed to indicate if they had seen a phosphene by pressing the left (‘no I did not see a phosphene’) or right (‘yes I did see a phosphene’) shift keys. After the REPT procedure the experimenter asked the participant to report which quadrants the participant had seen the phosphenes in to ensure they were the same as the initial testing.

### Transcranial direct current stimulation

tDCS was delivered by a battery driven portable stimulator (Neuroconn, Ilmenau, Germany) using a pair of 6 x 3.5 cm rubber electrodes in two saline soaked sponges.

Four different montages were used across the different experiments. In experiment 1 the active electrode was placed over Oz while the reference electrode was placed over the midline supraorbital area (see Fig.2B). In experiment 2 & 3 and the phosphene control experiment the active electrode was placed over Oz and the reference electrode was placed on the right cheek (see Fig.2C). In experiment 4 the active electrode was placed between F3 and Fz while the reference electrode was placed over the right cheek (see Fig.3E). In experiment 5 the electrodes were placed over Oz and between F3 and Fz.

In experiments 1, 2 & 4 each participant received both anodal and cathodal stimulations for a total of thirty minutes (fifteen minutes anodal, fifteen minutes cathodal) in two separate experimental sessions separated by a washout period of at least 24 hours, the order of which was randomized and counterbalanced across participants.

In experiment 3 each participant received both anodal and cathodal stimulations for a total of thirty minutes (fifteen minutes anodal, fifteen minutes cathodal) in two separate experimental sessions separated by a washout period of at least 24 hours, the order of which was randomized and counterbalanced across participants. These participants also received sham stimulation on one of the 3 days of testing, during the sham stimulation the machine ramped on for 5 seconds and switched off after 30 seconds of stimulation (ramping off over 5 seconds).

In experiments 5 each participant received both anodal-occipital + cathodal-prefrontal, and cathodal-occipital + anodal-prefrontal stimulations for a total of thirty minutes (fifteen minutes anodal, fifteen minutes cathodal) in two separate experimental sessions separated by a washout period of at least 24 hours, the order of which was randomized and counterbalanced across participants. These participants also received sham stimulation on one of the 3 days of testing, during the sham stimulation the machine ramped on for 5 seconds and switched off after 30 seconds of stimulation (ramping off over 5 seconds).

The experimenter was not blind to which polarity condition the participant was in from day to day. In experiment 1 the intensity used for stimulation was 1 mA, for all other experiments 1.5 mA was used. In the control tDCS modulation of phosphene thresholds experiment, the tDCS parameters were the same as in experiment 2. The intensity of the stimulation was set to 1.5mA and the active electrode placed over Oz and the reference electrode on the right cheek (see main text methods section and Fig. 2.C.)

### Statistical analysis of tDCS and TMS data

All correlational analysis, ANOVA’s and t-tests were run in SPSS (IBM, Armonk), and the LME’s were run in R (R Core Team, 2018) using the lme4 package(*68*).

For all linear mixed effects models with cathodal + anodal stimulation conditions, tDCS polarity (cathodal and anodal), block (D1, D2, P1, P2 – see S4A for timeline), and order of stimulation (cathodal stimulation first or second) were entered (without interaction terms) into the model as fixed effects. As random effects intercepts for subjects were entered into the model. P-values were obtained by likelihood ratio tests of the full model with tDCS polarity included versus the model without tDCS included.

For all linear mixed effects models with cathodal + anodal + sham stimulation conditions, tDCS polarity (cathodal, anodal, sham), block (D1, D2, P1, P2 – see S4A for timeline), and order of stimulation (cathodal stimulation first, second or third) were entered (without interaction terms) into the model as fixed effects. As random effects intercepts for subjects were entered into the model. P-values were obtained by likelihood ratio tests of the full model with tDCS polarity included versus the model without tDCS included.

## Acknowledgements

**General:** We thank Wolf Singer for his support in the project and for his helpful comments on the manuscript. We would also like to thank Roger Koenig for helpful comments.

## Author Contributions

JB collected and analyzed all of the MRI data. RK collected and analyzed all of the tDCS and TMS data. RK created all of the figures and RK, JB and JP wrote the manuscript.

## SUPPLEMENTARY MATERIAL

**Cortical excitability controls the strength of mental imagery**

- Supplementary Results
- Supplementary Figures (6 Figures)
- Supplementary Tables (3 Tables)

### Mock difference scores tDCS

#### 1 mA Occipital Stimulation

There was no significant main effect of tDCS polarity for the percentage change in mock priming (F(1,15) = 2.91, *p* = .11) or block on mock priming (Greenhouse-Geisser correction for violation of sphericity, F(1.90,28.51) = 1.69, *p* = .20), and no interaction between the two (Greenhouse-Geisser correction for violation of sphericity, F(1.68, 25.22) = .06, *p* = .92). When averaging across all of the blocks neither the cathodal or anodal condition mock priming was significantly different to 0 (t(15) 1.81, *p* = .10 and t(15) -.10, *p* = .92, respectively).

#### 1.5 Occipital Stimulation

There was no significant main effect of tDCS polarity for the percentage change in mock priming (F(1,15) = .07, *p* = .80) or block on mock priming (Greenhouse-Geisser correction for violation of sphericity, F(1.66, 24.97) = 1.23, *p* = .30), and no interaction between the two (Greenhouse-Geisser correction for violation of sphericity, F(1.59, 23.87) = 1.03, *p* = .36). When averaging across all of the blocks neither the cathodal or anodal condition mock priming was significantly different to 0 (t(15) 1.02, *p* = .33 and t(15) .66, *p* = .52, respectively).

#### 1.5 Occipital Stimulation (including sham)

There was no significant effect of tDCS polarity for the percentage change in mock priming (Mixed-effects analysis due to attrition Greenhouse-Geisser correction for violation of sphericity,: F(1.99, 28.82) = 1.23, *p* = .31) or block on mock priming (Mixed-effects analysis due to attrition Greenhouse-Geisser correction for violation of sphericity: F(2.12, 31.80) = .60, *p* = .56) and no interaction between the two (Mixed-effects analysis due to attrition Greenhouse-Geisser correction for violation of sphericity: F(3.40, 46.49) = .28, *p* = .87). When averaging across all of the blocks none of the polarity conditions were significantly different to 0 (sham: t(13) 1.80, *p* = .09, cathodal: t(15) .16, *p* = .87, and anodal: t(15) = .70, *p* = .50).

#### 1.5 Prefrontal Stimulation

There was no main effect of tDCS polarity for the percentage change in mock priming (F(1,15) = .02, *p* = .89) or block on mock priming (Greenhouse-Geisser correction for violation of sphericity, F(1.81, 27.20) = 1.18, *p* = .32), and no interaction between the two (Greenhouse-Geisser correction for violation of sphericity, F(1.78, 26.73) = 1.73, *p* = .20). When averaging across all of the blocks neither the cathodal or anodal condition mock priming was significantly different to 0 (t(15) -.18, *p* = .86 and t(15) -.59, *p* = .56, respectively).

#### 1.5 Prefrontal + Occipital Stimulation (including sham)

There was no significant effect of tDCS polarity for the percentage change in mock priming (Mixed-effects analysis due to attrition Greenhouse-Geisser correction for violation of sphericity,: F(1.06, 19.08) = .53, *p* = .76) or block on mock priming (Mixed-effects analysis due to attrition Greenhouse-Geisser correction for violation of sphericity: F(1.812, 32.62) = .60, *p* = .42) and no interaction between the two (Mixed-effects analysis due to attrition Greenhouse-Geisser correction for violation of sphericity: F(2.75, 42.11) = .46, *p* = .56). When averaging across all of the blocks none of the polarity conditions were significantly different to 0 (sham: t(17) .59, *p* = .55, cathodal occipital + anodal pFC: t(15) .23, *p* = .82, and anodal occipital + cathodal pFC: t(15) = .58, *p* = .57).

## Supplementary Figures

**Supplementary Figure S1.**
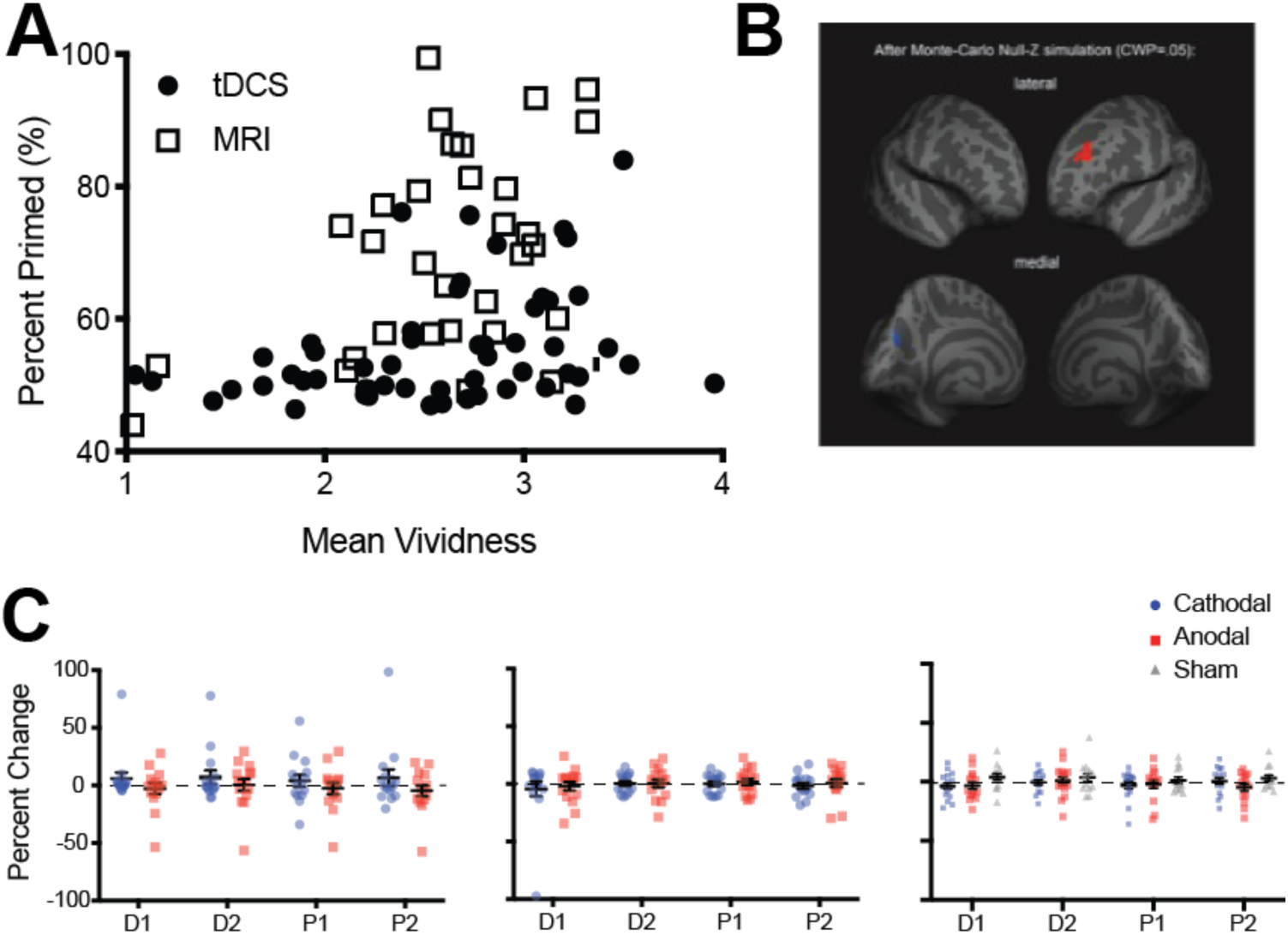
Imagery Vividness results. A. Data shows the correlation between mean vividness ratings (x-axis) and visual imagery priming (y-axis) for participants from both the MRI and tDCS experiments (tDCS experiment 2, 3 and 4). All possible participants’ data was included for the tDCS analysis (resulting in the analysis of 54 participant’s data). Only the correlation in the tDCS sample was significant, (r_s_ = .3655, p = .0033, Spearman’s rank-order correlation was used due to a violation of normality). The effect size was too small for significant results in the fMRI sample (r_s_ = .34, p = .065, Spearman’s rank-order correlation was used due to a violation of normality, N = 31). B. Surface-based whole brain analysis of the fMRI resting-state data: associations with subjective vividness. Corrected clusters showing associations with individual subjective vividness at a cluster-wise probability threshold of P < .05 (see also see Supplemental Table S3). The upper row shows a lateral view of the two hemispheres from an anterior perspective, whereas the lower row shows a medial view of them from the back. Multiple comparison correction was done using Monte Carlo Null-Z simulation (mc-z). No smoothing of the functional data was applied. Only two fMRI mean intensity clusters showed associations with subjective vividness that survived the correction for multiple comparisons: One cluster in the left rostralmiddlefrontal cortex showed a positive association (orange), and one smaller cluster in the left cuneus showed a negative association (blue). Note the similarity in the subjective vividness results with the ones in Bergmann et al. (2015), where only a volume cluster in left frontal cortex also showed a positive association with subjective vividness. Apparently, the positive relationship of subjective vividness with the anatomy of left frontal cortex is also reflected in the fMRI mean intensity levels of this region. C. tDCS effect on mean vividness ratings. Left panel: Occipital (1.5ma) – Vividness ratings were included in this experiment, which allowed us to look at subjective changes in imagery vividness that occur with changes in cortical excitability of the visual cortex. Red dots show the effect of anodal stimulation (increasing excitability) while blue dots show cathodal stimulation (decreasing excitability). Each dot represents an individual participant (one participant’s data is excluded from this analysis due to incorrect button presses on one of the days of testing, N =15). All data show means and error bars represent ±SEM’s. The data was again analyzed using percentage changed. We found no significant differences in the reported mean vividness of the imagined patterns (main effect of tDCS polarity: F(1,14) = 1.97, p = .18, main effect of block: F(3,42) = .73, p = .54, interaction: F(3,42) = .59, p = .63). Middle panel: tDCS of prefrontal cortex and mean vividness ratings. As vividness ratings were included in this experiment, we could also look at the subjective changes in imagery vividness that occur with changes in cortical excitability of the prefrontal cortex. Red dots show the effect of anodal stimulation (increasing excitability) while blue dots show cathodal stimulation (decreasing excitability). All data show means and error bars represent ±SEM’s. We again analyzed the data using percent change. There were no differences in the mean vividness ratings for either polarity of the tDCS (main effect: F(1,15) = .28, p = .61) or the block (main effect: F(3,45) = .77, p = .51), and there was no interaction between the two (F(3,45) = .07, p = .98). Right panel: tDCS of occipital cortex (1.5ma) – cathodal, anodal and sham stimulation, one participants data is removed due to incorrect button presses for vividness ratings. Red dots show the effect of anodal stimulation (increasing excitability) while blue dots show cathodal stimulation (decreasing excitability), and grey triangles show the effect of sham stimualtion. All data show means and error bars represent ±SEM’s. We again analyzed the data using percent change. There were no differences in the mean vividness ratings for any of the polarity conditions (Mixed effects analysis due to attrition : Polarity effect: F(1.95, 31.19) = 1.38, p = .27), there was also no effect of block (F(2.19, 34.98) = .94, p = .41) and no interaction (F(2.63, 36.82) = .98, p = .41).

**Supplementary figure S2.**
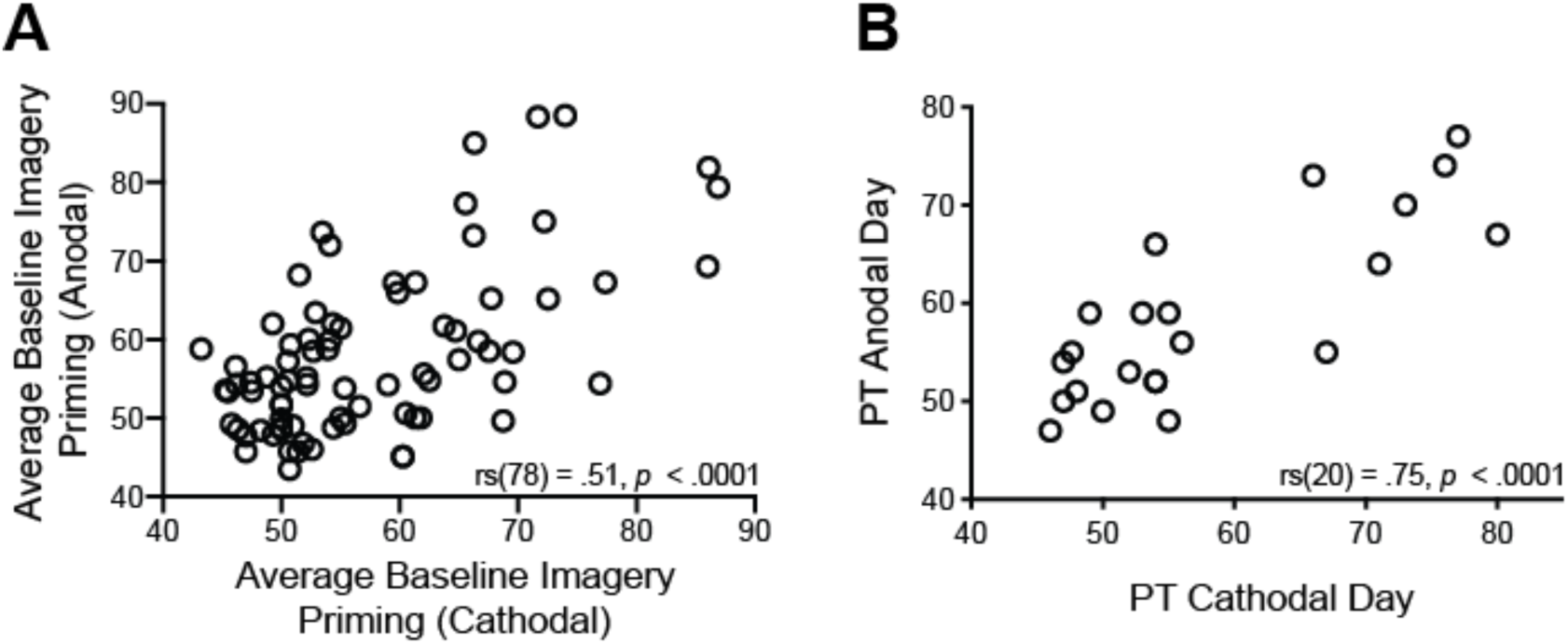
Re-test reliability for imagery strength **(A)** and Phosphene Thresholds **(B). A.** Scatterplot shows participants’ imagery strength measured by percent of binocular rivalry displays primed before tDCS stimulation across two days of testing (pre anodal and pre cathodal stimulation). Each data point represents one participant, 79 pairs in total. **B.** Scatterplot shows participants’ 60 percent phosphene thresholds (PT) before tDCS stimulation across the two days of testing. Each data point represents one participant, 21 pairs in total.

**Supplementary figure S3.**
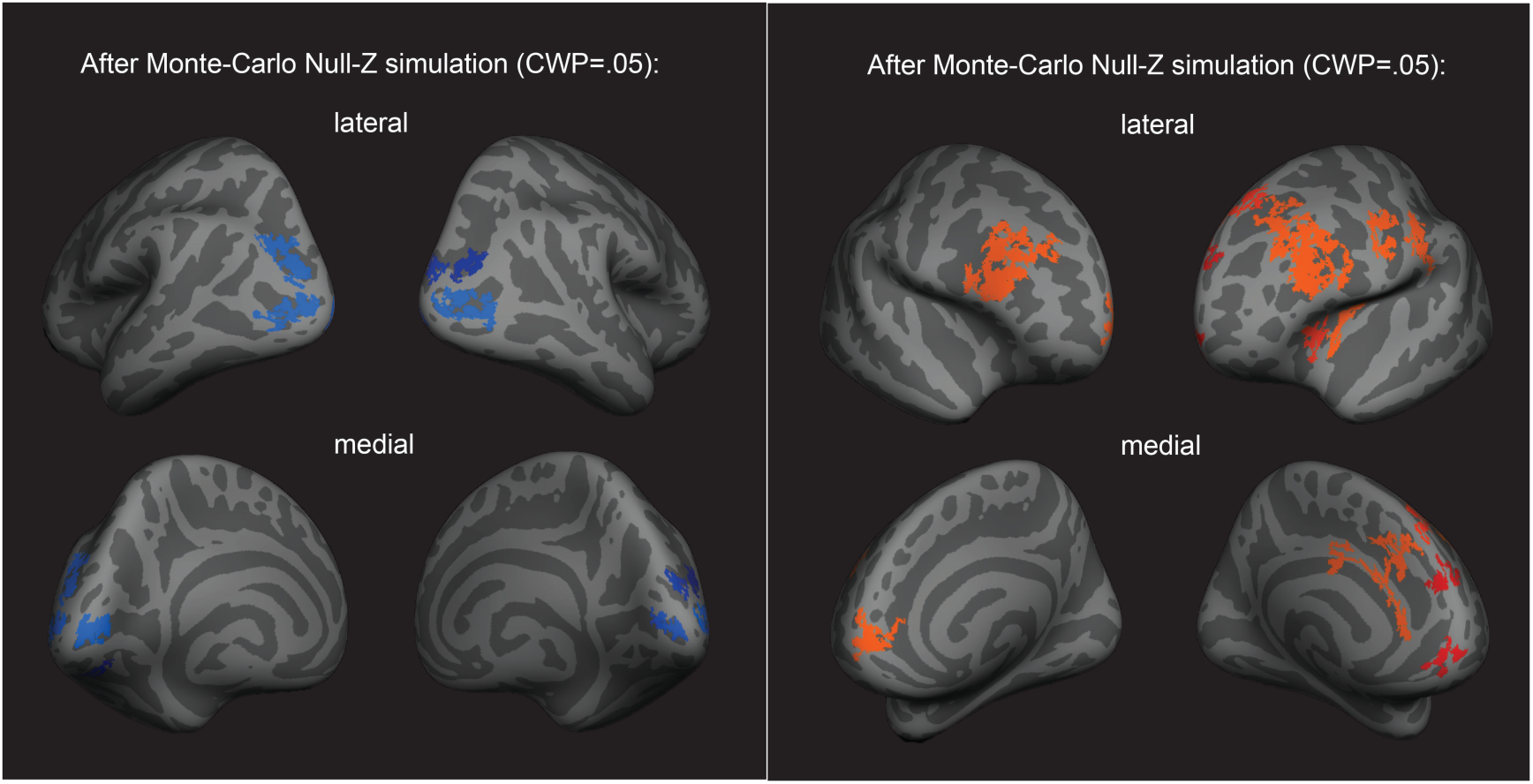
Left panel: Surface-based whole brain analysis of the fMRI resting-state data: negative associations with imagery strength. Corrected clusters showing a *negative* association with individual imagery strength at a cluster-wise probability threshold of *P* < .05 (also see Supplementary Table S1). In both rows, the two hemispheres are shown from the back. Multiple comparison correction was done using Monte Carlo Null-Z simulation (mc-z). Right panel: Surface-based brain analysis of the fMRI resting-state data and imagery: positive associations with imagery strength. Corrected clusters showing a *positive* association with individual imagery strength at a cluster-wise probability threshold of *P* < .05 (also see Supplementary Table S2). In both the lateral and medial view, the hemispheres are shown from the front. Multiple comparison correction was done using Monte Carlo Null-Z simulation (mc-z). No smoothing of the functional data was applied. In line with the correlation analyses using normalised fMRI mean intensity of atlas- and retinotopically defined areas, only fMRI mean intensity clusters in frontal areas showed positive associations with imagery strength (% primed).

**Supplementary figure S4.**
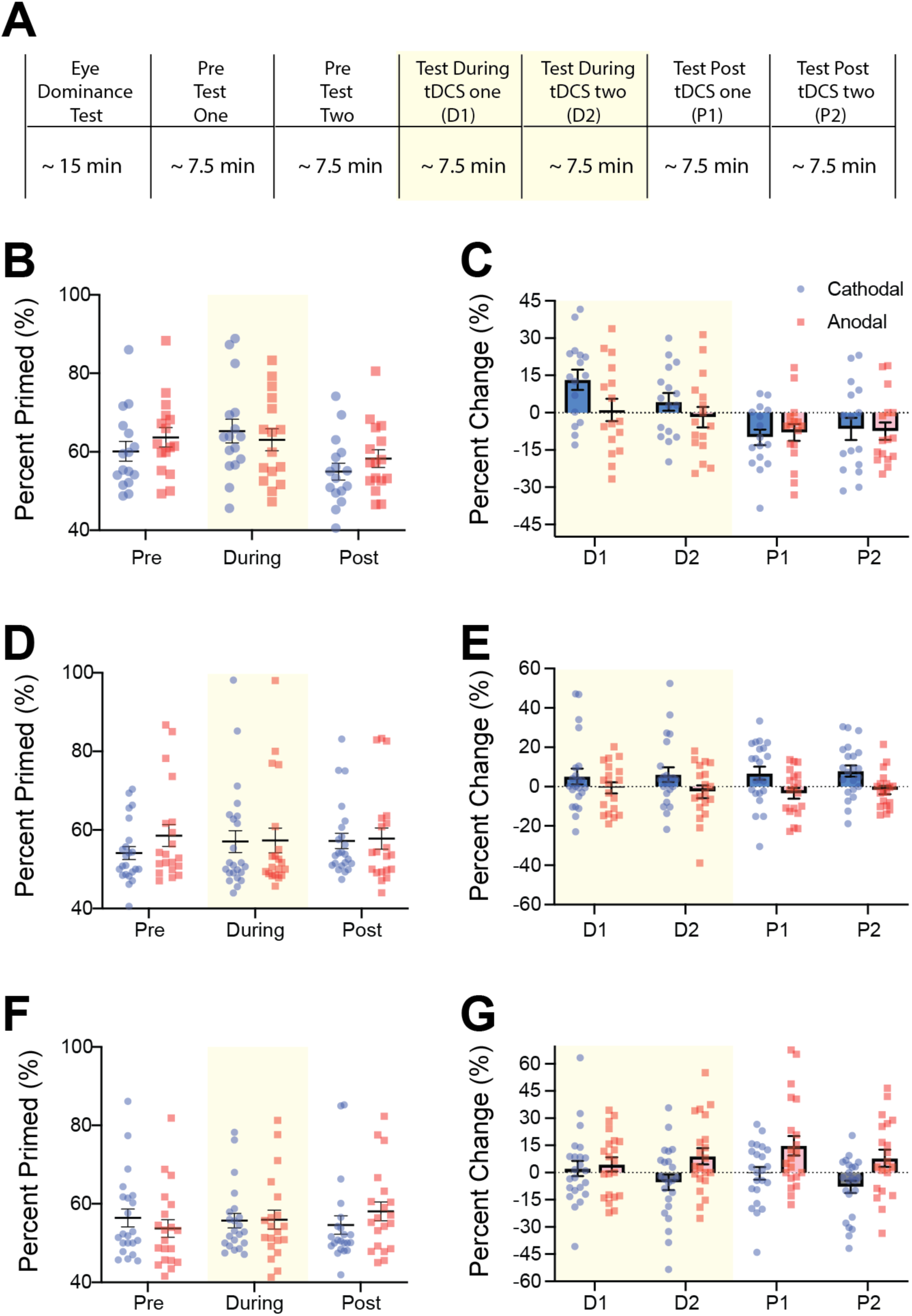
**A**. Experimental timeline for all tDCS experiments. Spread of individual data points for raw data and difference scores for experiment 1 (1mA Occipital: **B & C**), experiment 2 (1.5mA Occipital: **D & E**) and experiment 4 (1.5ms PreFronal: **F & G**). Blue data points represent individual subjects’ cathodal stimulation changes while red data points represent anodal stimulation changes. For the raw data (**B, D & F**) the figure shows data averaged across pre, during and post stimulation blocks. For the difference scores data (**C, E & G**) the figure shows difference scores for each imagery block, two during (D1 and D2, shaded yellow area) and two after tDCS (P1 and P2). All error bars show ±SEMs.

**Supplementary figure S5.**
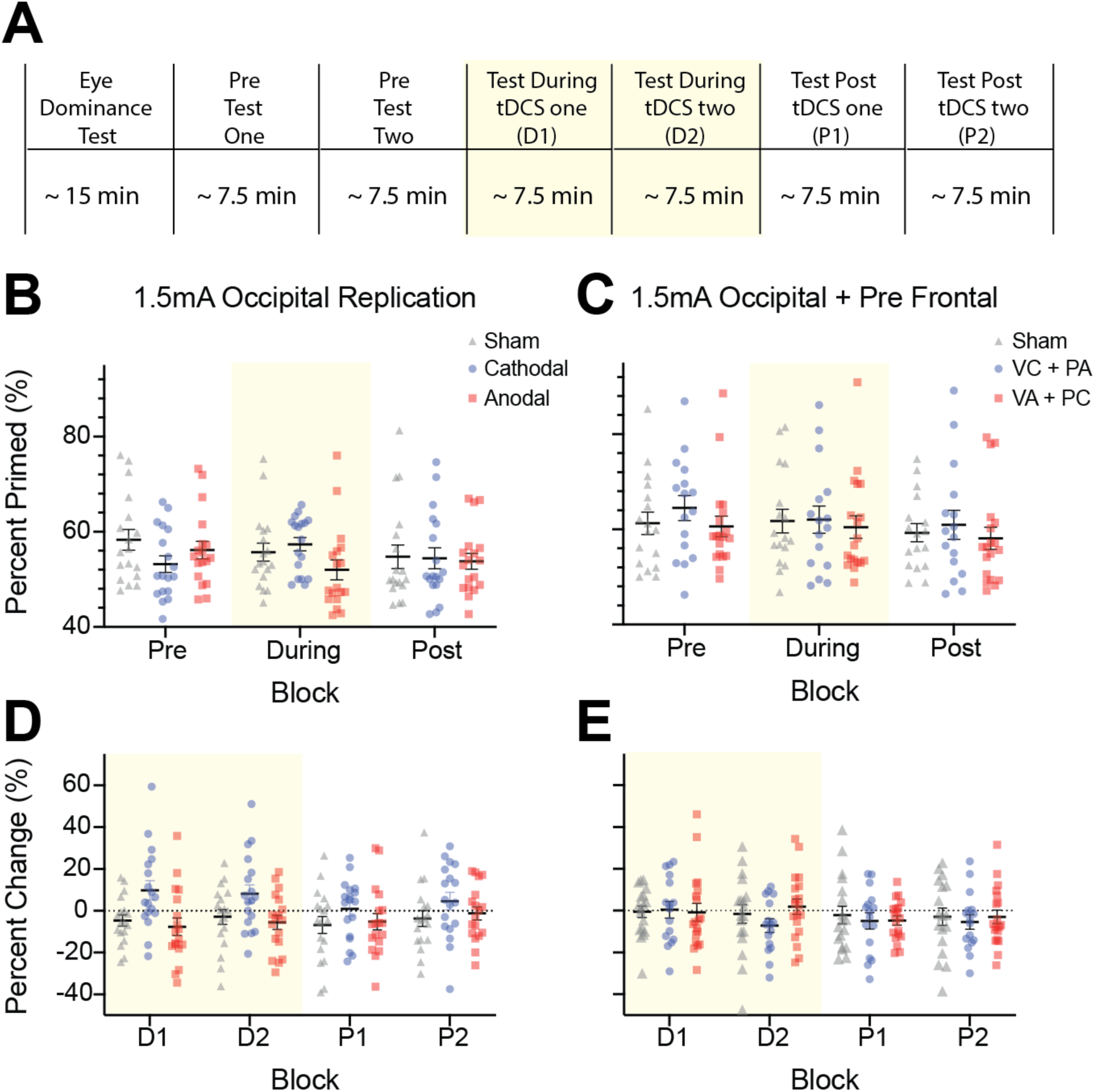
***A.*** Experimental timeline for all tDCS experiments. Spread of individual data points for raw data and difference scores for experiment 3 (Occipital stimulation: **B & D**), experiment 5 (Occipital + Prefrontal stimulation: **C & E**). For **B** and **C** blue data points represent individual subjects’ cathodal stimulation changes, red data points represent anodal stimulation changes and grey data points represent sham stimulation. For **D & E** blue data points represent cathodal-occipital and anodal-prefrontal stimulation, while blue represents anodal-occipital and cathodal-prefrontal stimulation, and grey data points represents sham stimulation. For the raw data (**B & C**) the figure shows data averaged across pre, during and post stimulation blocks. For the difference scores (percent change) data (**D & E**) the figure shows difference scores (percent change) for each imagery block, two during (D1 and D2, shaded yellow area) and two after tDCS (P1 and P2). All error bars show ±SEMs.

**Supplementary figure S6.**
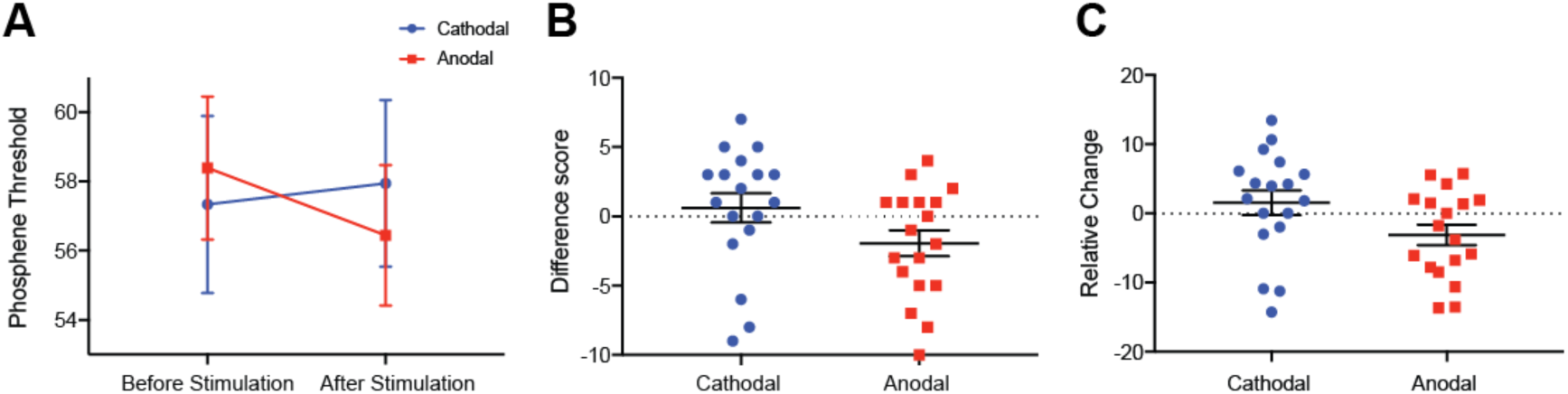
tDCS modulation of phosphene thresholds. **A.** Data shows phosphene thresholds (PT) before cathodal (left hand side, blue data points) and before anodal (left hand side, red data points) and after cathodal (right hand side, blue data points) and after anodal stimulation (right hand side, red data points). A significant interaction between tDCS polarity and PT session was found (F(1,17) = 6.16, *p* = .02). **B.** We then looked at the difference scores for each participant’s phosphene thresholds in the cathodal and anodal conditions. This difference score was calculated with the following equation: PT_(after tDCS)_ – PT_(before tDCS)_. Data shows participants’ phosphene threshold differences scores with positive scores indicating that PTs have increased after tDCS (in the cathodal condition, blue bar) while negative scores indicate that PTs have decreased after tDCS (in the anodal condition, red bar). There was a significant difference between the anodal and cathodal conditions with anodal PT changes being significantly lower than cathodal (t(17) = 2.48, *p* = .02). **C**. To assess whether or not some participants were driving these results, e.g. it might be that participants with high phophene thresholds are driving the results, relative difference scores were also calculated: ((PT_(after tDCS)_ – PT(_before tDCS))_/ PT_(before tDCS)_)*100. Using this method of analysis the same pattern of results was found with cathodal stimulation increasing phosphene thresholds in comparison to anodal stimulation (t(17) = 2.70, *p* = .015). All error bars show ±SEMs.

## Supplementary Tables

**Supplementary Table S1.** Surface-based whole brain analysis of the fMRI resting-state data: Corrected clusters showing a significantly *negative* association with individual imagery strength at a cluster-wise probability threshold of *P* < .05 (also see Supplementary Fig. S1).

| Cluster No. | Maximum voxel-wise significance in cluster | Peak vertex annotation | Size (mm<sup>2</sup>) | MNI 305 coordinates of peak vertex | | | Cluster-wise $P$-value | NVtxs | Laterality | Area label |
| --- | --- | --- | --- | --- | --- | --- | --- | --- | --- | --- |
| 1 | -4.613 | 11477 | 552.22 | -24.7 | -86.6 | 12.8 | 0.0002 | 903 | L | lateraloccipital |
| 2 | -4.172 | 97048 | 535.7 | -20.6 | -96.9 | 3.3 | 0.0002 | 734 | L | lateraloccipital |
| 3 | -3.936 | 130756 | 534.22 | -13.3 | -86.4 | 5.3 | 0.0002 | 698 | L | pericalcarine |
| 4 | -3.053 | 63050 | 248.57 | -19.5 | -80.4 | -7.3 | 0.0024 | 298 | L | lingual |
| 5 | -2.848 | 162967 | 203.8 | -31.3 | -59.6 | -8.4 | 0.01276 | 374 | L | fusiform |
| 1 | -4.488 | 69964 | 301.97 | 17 | -96.6 | 0.8 | 0.0004 | 408 | R | pericalcarine |
| 2 | -4.021 | 71800 | 560.64 | 27.4 | -92.3 | 1.2 | 0.0002 | 785 | R | lateraloccipital |
| 3 | -3.739 | 117794 | 239.69 | 14.6 | -91.3 | 19 | 0.0028 | 301 | R | lateraloccipital |
| 4 | -2.502 | 104614 | 214.22 | 35.9 | -82.5 | 18.4 | 0.00519 | 366 | R | inferiorparietal |

Note: Correction for multiple comparisons at a cluster-wise probability threshold of $p < .05$ was done using a pre-cached Monte Carlo Null-Z simulation with 10000 iterations. The locations of the clusters strongly overlap with the locations of the atlas- and retinotopically defined areas that showed a negative association with imagery strength. Abbreviations: L = left hemisphere, R = right hemisphere; NVtxs = Number of vertices.

**Supplementary Table S2.**
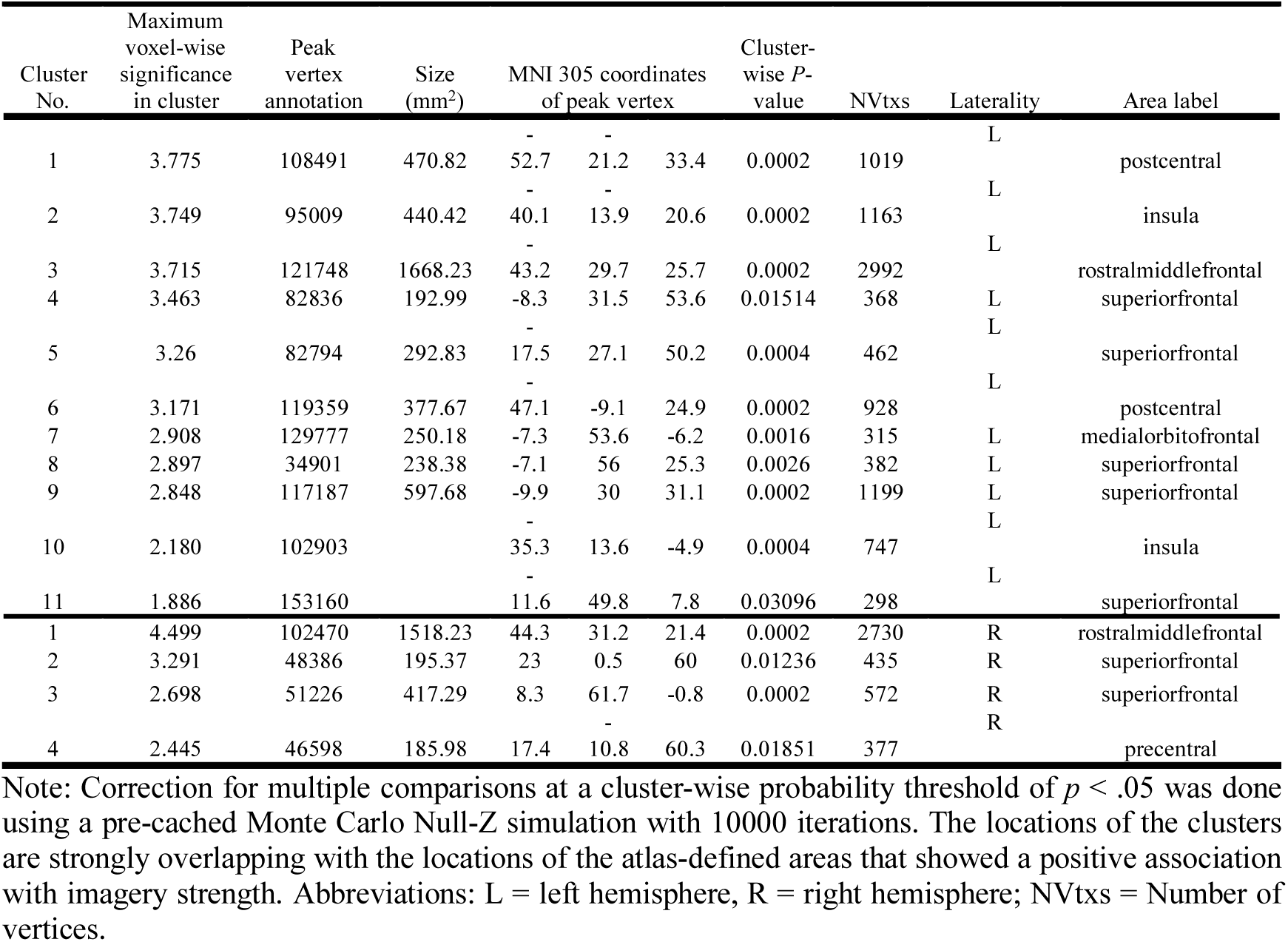
Surface-based whole brain analysis of the fMRI resting-state data: Corrected clusters showing a significantly *positive* association with individual imagery strength at a cluster-wise probability threshold of *P* < .05 (also see Supplementary Fig. S2).

**Supplementary Table S3.**
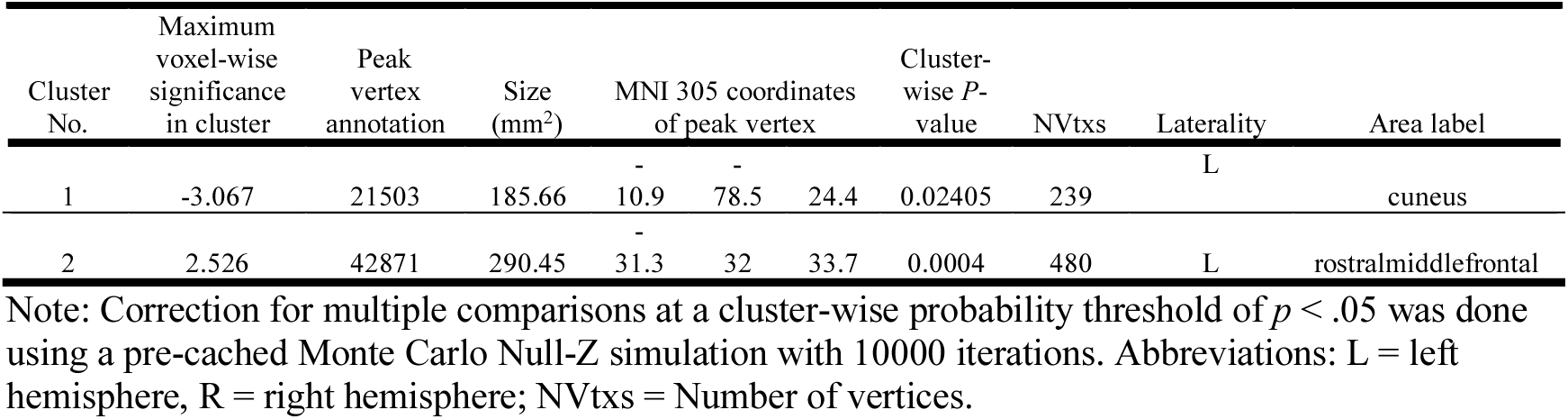
Surface-based whole brain analysis of the fMRI resting-state data: Corrected clusters showing significantly positive and negative associations with individual subjective vividness at a cluster-wise probability threshold of *P* < .05 (also see Supplementary Fig. S3).

